# The human developing cerebral cortex is characterized by an increased de novo expression of lncRNAs in excitatory neurons

**DOI:** 10.1101/2023.10.26.564246

**Authors:** David A. Morales-Vicente, Ana C. Tahira, Daisy Woellner-Santos, Murilo S. Amaral, Maria G. Berzoti-Coelho, Sergio Verjovski-Almeida

## Abstract

**Background:** Outstanding human cognitive abilities are computed in the cerebral cortex, a mammalian-specific brain region and the place of massive biological innovation. Long noncoding RNAs (lncRNAs) have emerged as gene regulatory elements with higher evolutionary turnover than mRNAs. The many lncRNAs identified in neural tissues make them candidates for molecular sources of cerebral cortex evolution and disease. Here, we characterized the genomic and cellular shifts that occurred during the evolution of the lncRNA repertoire expressed in the developing cerebral cortex of humans and explored their role in the evolution of this brain region.

**Results:** Using systems biology approaches and comparative transcriptomics, we comprehensively annotated the cortical transcriptomes of humans, macaques, mice, and chickens and classified human cortical lncRNAs into evolutionary groups as a function of their predicted minimal ages. LncRNA evolutionary groups showed differences in expression levels, splicing efficiencies, transposable element contents, genomic distributions, and transcription factor binding to their promoters. Furthermore, older lncRNAs showed preferential expression in germinative zones, outer radial glial cells, and cortical inhibitory neurons. In comparison, younger lncRNAs showed preferential expression in cortical excitatory neurons, belonged to human-specific gene coexpression modules, and were dysregulated in autism spectrum disorder.

**Conclusions:** These results suggest a shift in the roles of cortical lncRNAs over evolution, highlighting the antique lncRNAs as a source of molecular evolution of conserved developmental programs; conversely, the *de novo* expression of primate and human-specific lncRNAs are sources of molecular evolution and dysfunction of cortical excitatory neurons.

## Background

The cerebral cortex is a primary information processing center of the central nervous system, is key to the evolution of higher cognition, and is affected in neurodevelopmental disorders. It comprises billions of excitatory projection neurons (glutamatergic) and inhibitory interneurons (GABAergic) assembled in local circuits intertwined with glial and vascular cells arranged in a six-layered architecture on the outer surface of the mammalian brain [1, 2]. The cerebral cortex evolved from the dorsal pallium after the divergence of mammals and sauropsids (reptiles and birds) approximately 300 million years ago (MYA), and it is endowed with incredible plasticity, evident in the diverse neocortical sizes and shapes [3, 4]. Primates present an expanded brain with more total neurons than most mammalian species. The human cerebral cortex has further expanded, differentiating us from our closest living relatives. These expansions and increased diversity of neuronal cell types are likely responsible for the increased computational capacity and unparalleled human cognition [5].

At the cellular level, the human developing cerebral cortex presents an augmented proliferative capacity of neural progenitors (radial glial cells, RGCs), especially from the outer subventricular zone (outer radial glial cells, oRGCs) [4], as well as an improvement in the information processing capability of mature excitatory neurons [5]. Understanding the molecular basis of these differences is critical to unveiling the evolution of human higher cognition and having a deeper comprehension of how they are disrupted in diseases [3]. In this line, a considerable effort has been made in the past decade to identify those changes, finding that duplications of protein-coding genes and modifications in gene-regulatory regions have altered the transcriptional landscape of different cell populations in the developing cerebral cortex [1, 6]. This extensive work has mainly focused on changes in the expression of protein-coding genes; expanding this analysis to the human noncoding transcriptome is crucial to improve our understanding of the gene-regulatory modifications that have led to the evolution of the human cerebral cortex.

Long noncoding RNAs (lncRNAs) are noncoding genes transcribed into RNAs longer than 200 nucleotides that do not translate into functional proteins [7]. This heterogeneous group of lncRNAs is transcribed by RNA pol II and shares molecular features with mRNAs, such as being 5’ capped, spliced, and polyadenylated; despite the molecular similarities with mRNAs, lncRNAs also present features that differentiate them, including higher tissue specificity compared to mRNAs, distinct chromatin modifications at the promoter region, cell nucleus enrichment, inefficient splicing, and less stability than mRNAs [8, 9]. Although these features may point to lncRNAs as mere transcriptional noise, it has been shown that at least a fraction of lncRNAs or the act of their transcription have gene-regulatory functions [9]. Interestingly, after testes, neural tissues express the most significant number of lncRNAs in tetrapods [10–12], and several lncRNAs have been characterized as functional regulatory RNAs of different stages of corticogenesis [13].

Unlike protein-coding genes that have evolved mainly by gene duplications, lncRNAs have preferentially evolved by *de novo* expression and exonization mediated by transposable elements (TEs) [14]. The *de novo* expression and the significant contributions of TEs to the evolution of lncRNAs explain the reduced constraint under which lncRNAs evolved compared to protein-coding genes [14]. Furthermore, the first highly evolving human-specific region (HAR, human accelerated region) was identified inside the lncRNA *HAR1F*, which is expressed in the developing cerebral cortex [15]. In addition, it has been shown in mammals that lncRNAs are a source of cellular plasticity due to their capacity to acquire new functional modalities [16], and some lncRNAs are a source of newly identified peptides [17]. This faster evolutionary turnover of lncRNAs compared to mRNAs and their gene-regulatory functions make lncRNAs good candidates for molecular drivers of biological innovations, especially in the cerebral cortex.

Here, we used systems biology approaches and comparative transcriptomics to characterize the evolution of the lncRNA repertoire of the developing cerebral cortex. Our analyses identified signatures of a functional shift of cortical lncRNAs throughout evolution and pointed to lncRNAs as a source of molecular innovation of the human cerebral cortex.

## Results

### Assemblies of new comprehensive transcriptomes improve the annotation of lncRNAs in humans and other three vertebrate model organisms

To better understand the relevance of changes in the lncRNA repertoire during the evolution of the cerebral cortex, we first needed to characterize the evolution of human lncRNAs. After the testis, the vertebrate nervous system contains more specific lncRNAs than other body tissues [10–12]. Consequently, the lack of extensive annotation of RNA-seq libraries from the prenatal cerebral cortex has hampered the reconstruction of more representative lncRNA gene models for this brain region. To avoid misidentifying homologous lncRNAs due to the differences in the completeness of the human, rhesus macaque, mouse, and chicken transcriptome annotations, we generated and annotated new comprehensive catalogs of lncRNAs.

We gathered an extensive collection of RNA-seq libraries encompassing different stages of cerebral cortex and cerebral pallium development. This collection of RNA-seq libraries includes recently published datasets from humans, rhesus macaques, and mice [12, 18, 19] (Additional file 1: Fig. S1a–c and Additional file 2: Table S1); additionally, we generated new bulk RNA-seq data from the chicken pallium (Additional file 1: Fig. S1d and Additional file 2: Table S1). We removed RNA-seq libraries with high 3’ bias (Additional file 1: Fig. S2a–d), and filtered libraries were used to assemble new transcriptome models for all species. We developed a genome-assisted approach which integrates efficient and more accurate bioinformatics tools that we applied to the collection of short-read RNA-seq libraries (see methods and Additional file 1: Fig. S3a). We used the latest available genomes as references (hg38, rheMac10 [20], mm39 [21], and galGal6 [22, 23]), most of which were based on long-read sequencing. In addition, we collected publicly available IsoSeq long reads from rhesus macaques and chickens (see methods and Additional file 1: Fig. S3b) to help improve the transcriptome assembly based on short reads. Large groups of lncRNAs were annotated in all transcriptome assemblies following stringent criteria (see methods and Additional file 1: Fig. S3c). To generate the final comprehensive catalog of lncRNAs of the four species, lncRNAs from public lncRNA databanks [10, 12, 24] were incorporated into the final transcriptomes.

Of these new transcriptomes, lncRNAs represent the category with more annotated genes (Additional file 1: Fig. S4a–d), corroborating the widespread expression of lncRNAs in vertebrates [10–12]. Our new assembly contributed over 40% of all lncRNAs in humans, rhesus macaques, and chickens (Additional file 1: Fig. S4e–h). Remarkably, we significantly improved the number of reads mapped to annotated features (Additional file 1: Fig. S4i–l). In addition to contributing to the annotation of new lncRNAs, we identified new protein-coding genes, pseudogenes, and other gene types, showing the robustness of our approach for identifying new coding and noncoding transcripts across different organisms, especially for species with poorly annotated transcriptomes, such as chickens and rhesus macaques.

### Syntenic conservation allows the classification of cortical lncRNAs into minimal evolutionary age (MA) groups

The fast evolutionary turnover of lncRNAs at a primary sequence level precludes the identification of orthologs in other species using conventional approaches, particularly for species that have diverged for long evolutionary periods. Syntenic conservation of genes (shared genomic positions between two species) has been used to improve the identification of homologous lncRNAs, allowing the discovery of syntenic conserved lncRNAs across vertebrates [10, 25]. To identify changes in the lncRNA repertoire throughout the evolution of the cerebral cortex, a bioinformatics pipeline was developed to systematically classify lncRNAs into evolutionary ancestry groups based on the syntenic conservation of human lncRNA genes in the rhesus macaque, mouse, and chicken genomes (Fig. 1a–b, see Methods). To reduce the chance of type I errors, we required that the set of syntenic lncRNAs identified when using the human lncRNA transcriptome as a query be the same when using the other species as a query, thus allowing the identification of lncRNAs with reciprocal syntenic signals (Fig. 1b). To improve the classification of human lncRNAs into minimal-evolutionary-age (MA) groups, syntenic conservation data from lncRNAs in public databases [10, 12, 26] were incorporated into the identified reciprocal syntenic conservation. The human lncRNAs identified as syntenic to chicken, mouse, and macaque lncRNAs were clustered into 300 million years ago (Mya), 90 Mya, and 25 Mya MA groups, respectively (Fig. 1c). LncRNAs that did not share synteny with any species using our approach or were not annotated as syntenic conserved in public databases were classified as human-specific lncRNAs (Additional file 3: Table S2). Other lncRNAs were classified as weakly syntenic and not further considered for subsequent analysis if they presented nonreciprocal syntenic conservation and were not annotated as syntenic conserved in public databases. The MA groups were clustered around landmarks of cerebral cortex evolution: before and after the evolution of the cerebral cortex from the dorsal pallium, throughout the expansion of the cerebral cortex in the primate lineage, and during the current evolution of the cerebral cortex in humans (Fig. 1c).

**Fig. 1.**
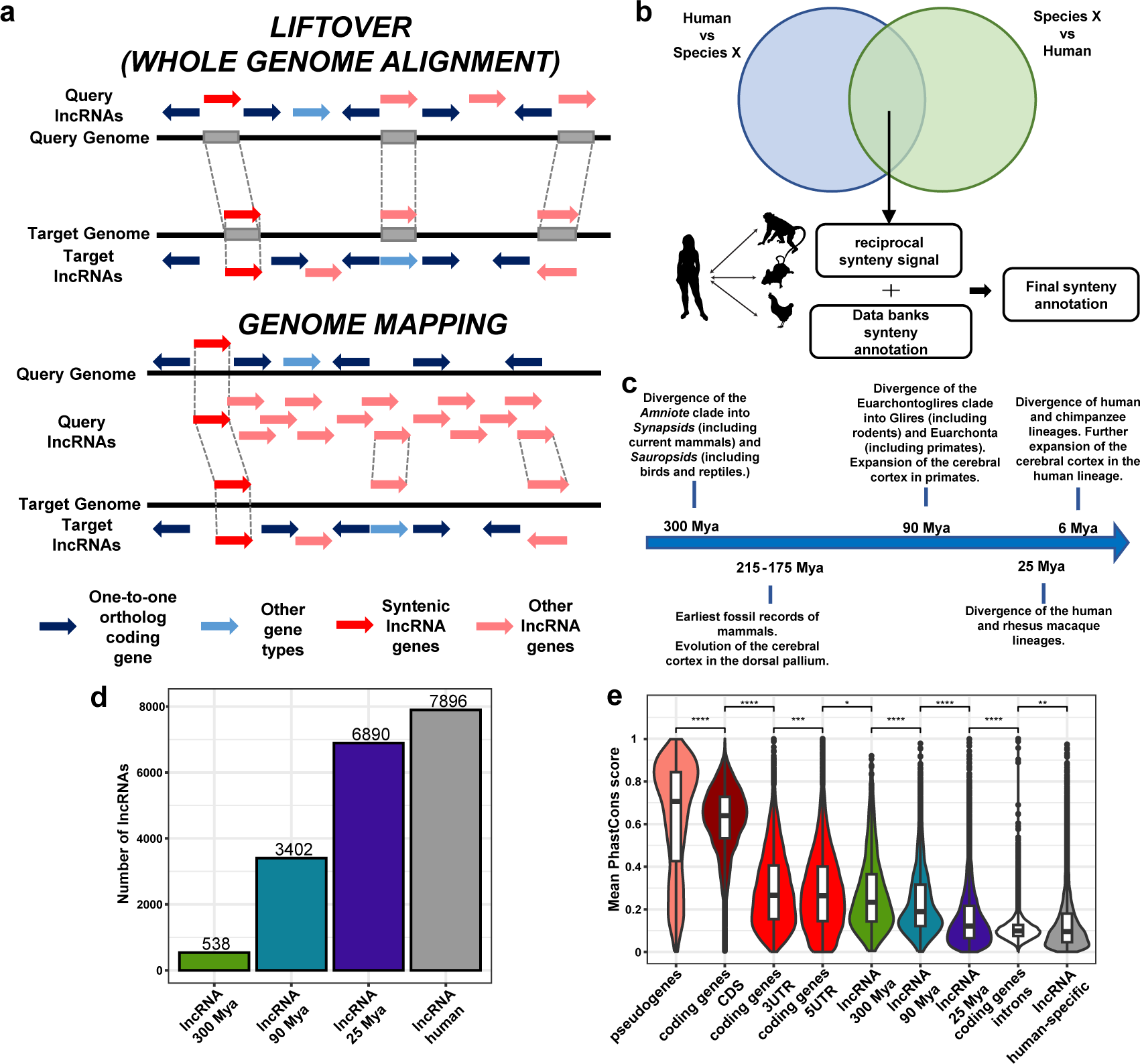
Classification of cortical lncRNAs into minimal evolutionary age groups. **a** Depiction of the approaches used to identify syntenic lncRNAs between two species. Top: whole genome alignment, bottom: genome mapping. **b** Workflow for the identification of reciprocal syntenic orthologs of human lncRNAs. **c** Landmarks of cerebral cortex evolution in the human lineage. **d** Distribution of the lncRNA genes in each minimal-evolutionary-age (MA) group category. **e** Mean PhastCons conservation scores of protein-coding genes, pseudogenes, and lncRNA MA groups. Statistics: All statistics are one-sided (greater) Wilcoxon tests. ns: p greater than 0.05, *: p equal to or less than 0.05, **: p equal to or less than 0.01, ***: p equal to or less than 0.001, ****: p equal to or less than 0.0001.

After classifying the human lncRNA repertoire into MA groups, we identified the set of genes expressed throughout the prenatal development of the cerebral cortex using a total of 189 bulk RNA-seq libraries from the PsychEncode project [18] and two public single-cell RNA-seq datasets from the developing cerebral cortex [27, 28]. Genes were identified as cortical genes if they were expressed with at least 0.5 TPM (Additional file 3: Table S2) in all samples considered at each developmental window/region pair (Table 1) or if they were detected as differentially expressed (DE) in one single-cell cluster (Additional file 4: Table S3). A final number of 40,348 genes were detected as expressed in the developing cerebral cortex, of which 18,726 cortical lncRNAs were classified into one of four MA groups (Fig. 1d and Additional file 3: Table S2). The number of cortical lncRNAs increases throughout the evolution of the human lineage, with only 2.8% of them identified as appearing before the evolution of the cerebral cortex and approximately 42% of them being specific to humans (Fig. 1d), in line with the fast evolutionary turnover described for lncRNAs [10, 11].

**Table 1.** Human cerebral cortex developmental window-region pairs.

| original brain region | PsychEncode developmental window |  |  |  |  |
| --- | --- | --- | --- | --- | --- |
|  | W1* | W2* | W3* | W4* | W5* |
| dorsolateral prefrontal cortex | 2 | 6 | 4 | 4 | 3 |
| medial prefrontal cortex | 2 | 6 | 4 | 4 | 4 |
| orbital prefrontal cortex | 2 | 6 | 2 | 3 | 4 |
| ventrolateral prefrontal cortex | 0 | 6 | 4 | 4 | 4 |
| primary motor-somatosensory cortex | 2 | 0 | 3 | 0 | 0 |
| primary motor (m1) cortex | 0 | 6 | 1 | 3 | 4 |
| primary somatosensory (s1) cortex | 0 | 6 | 0 | 3 | 4 |
| occipital neocortex | 2 | 0 | 0 | 0 | 0 |
| primary visual (v1) cortex | 0 | 6 | 4 | 3 | 4 |
| parietal cortex | 1 | 0 | 0 | 0 | 0 |
| posterior inferior parietal cortex | 0 | 6 | 4 | 3 | 4 |
| primary auditory (a1) cortex | 0 | 6 | 4 | 3 | 4 |
| inferior temporal cortex | 0 | 6 | 2 | 2 | 4 |
| superior temporal cortex | 0 | 4 | 4 | 3 | 4 |

\*Number of libraries per brain region (indicated at left) in each developmental window (Wn, indicated at top) that were used to identify the expression of lncRNAs in the developing cerebral cortex. From a total of 189 libraries (Additional file 2: Table S1).

Next, we checked the phastCons scores [29] to test the conservation status among MA groups of cortical lncRNAs. The conservation scores significantly decreased throughout the evolution of cortical lncRNAs (Fig. 1e), indicating that the approach implemented here correctly classified lncRNAs into sequential evolutionary groups. lncRNA genes were further classified depending on their position in the genome relative to protein-coding genes (Fig. 2a, left). The relative abundance of lncRNA types changed through evolution, with older lncRNAs being mostly antisense, while overlapping, intergenic proximal, and intronic lncRNA fractions increased in the younger lncRNAs (Fig. 2a, right). Significantly, the number of intronic lncRNAs has increased in the human lineage (Fig. 2a, right), as 80.9% of the total intronic lncRNAs are human specific.

**Fig. 2.**
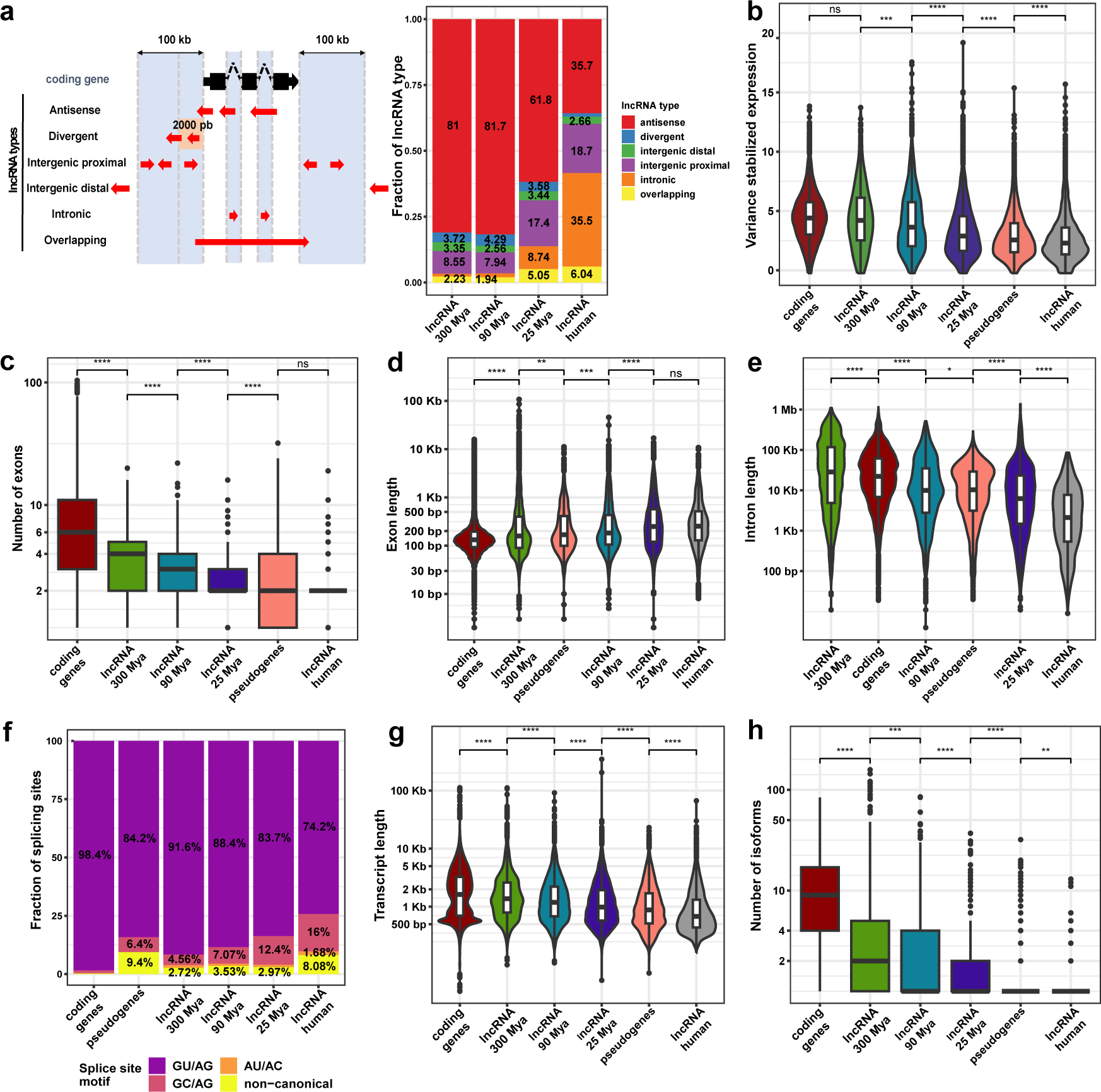
Differences in the genomic features of cortical lncRNAs. **a** Left, schematic depiction of the classification of lncRNAs based on their position regarding protein-coding genes; Right, distribution of lncRNA types among the lncRNA minimal-evolutionary-age (MA) groups. **b** Expression levels of protein-coding genes, pseudogenes, and lncRNA MA groups. **c** Number of exons distribution among protein-coding genes, pseudogenes, and lncRNA MA groups. **d–e** Like **c** but showing the exon and intron length distribution, respectively. **f** Frequency of splicing motifs among protein-coding genes, pseudogenes, and lncRNA MA groups. **g–h** Transcript length distribution and the number of isoforms among protein-coding genes, pseudogenes, and lncRNA MA groups. **Statistics:** All statistics are one-sided (greater) Wilcoxon tests. ns: p greater than 0.05, *: p equal to or less than 0.05, **: p equal to or less than 0.01, ***: p equal to or less than 0.001, ****: p equal to or less than 0.0001.

### Older lncRNAs have enhanced expression strength, splicing efficiency, and loci complexity

After classifying the cortical lncRNAs into the MA groups, we assessed the differences in their genomic features. We found that the MA groups follow an expression gradient, where older lncRNAs reach more robust expression levels in the human cortex than younger lncRNAs (Fig. 2b). Interestingly, the oldest group of cortical lncRNAs achieves similar expression levels to protein-coding genes (Fig. 2b), which shows that the general lower expression of lncRNAs compared to protein-coding genes [10, 11] is masked by the significant difference in gene expression levels between old and young lncRNAs.

We thought the documented differences between mRNA and lncRNA splicing efficiency [10] might also be masked by the different MA groups, as is the case for gene expression levels. Therefore, we assessed differences in splicing efficiency among the MA groups. The analyses were restricted to a set of expression and type-matched genes, as differences in expression levels and lncRNA type may interfere with the splicing differences among the MA groups. The following splicing features were evaluated: the number of exons, exon lengths, intron lengths, and splicing motif frequency. Similar to the gene expression levels, the number of exons of the lncRNA MA groups follows a gradient, where older lncRNA populations have significantly more exons on average than younger populations (Fig. 2c). Nevertheless, in contrast to expression levels, mRNAs present a considerably higher number of exons than all lncRNA MA groups (Fig. 2c). Simultaneously, older lncRNAs have significantly shorter exon lengths and longer intron lengths than younger lncRNAs (Fig. 2d–e), while mRNAs have, in general, shorter exon lengths than all lncRNA MA groups (Fig. 2d); interestingly, the oldest MA group has more extensive intron lengths than mRNAs (Fig. 2e). Furthermore, older lncRNAs present a higher frequency of stronger splicing motifs than younger lncRNAs (Fig. 2f). In summary, these features indicate an enhancement of splicing efficiency in older lncRNAs that suggests a gain in functionality during evolution.

It has been shown that longer transcripts have features of dynamic expression associated with lncRNA functionality [12]. To further evaluate a possible increase in the functionality of older lncRNAs, we assessed the length of the transcripts and the number of isoforms among the MA groups. Older lncRNAs have significantly longer transcripts and significantly more isoforms than younger lncRNAs (Fig. 2g–h), indicating an increase in locus complexity for older lncRNAs; however, all lncRNAs have reduced locus complexity compared with protein-coding genes (Fig. 2h).

### LncRNA evolutionary groups show distinct distributions of transposable element insertions but shared nuclear retention

Transposable elements (TEs) are the main drivers of lncRNA diversification and evolution [10, 14]; therefore, TEs might be involved in the differences in transcript complexity among lncRNA MA groups. To evaluate this scenario, the incidence of TE insertions was assessed. Interestingly, all lncRNAs show a similar percentage of TE occurrence (56.51-58.85%, Fig. 3a) that differs from the depletion of TE insertion in pseudogenes and protein-coding genes; nevertheless, 3’ UTRs present an increased incidence of TE insertions (Fig. 3a), indicating that this UTR with regulatory functions is tolerant to TE insertions. Nevertheless, the extent of the TE body inserted in genes differs among lncRNA MA groups. Older lncRNAs incorporated larger fractions (72.2%) of full-length TE sequences than younger lncRNAs (46.1 to 66.3%) (Fig. 3b).

**Fig. 3.**
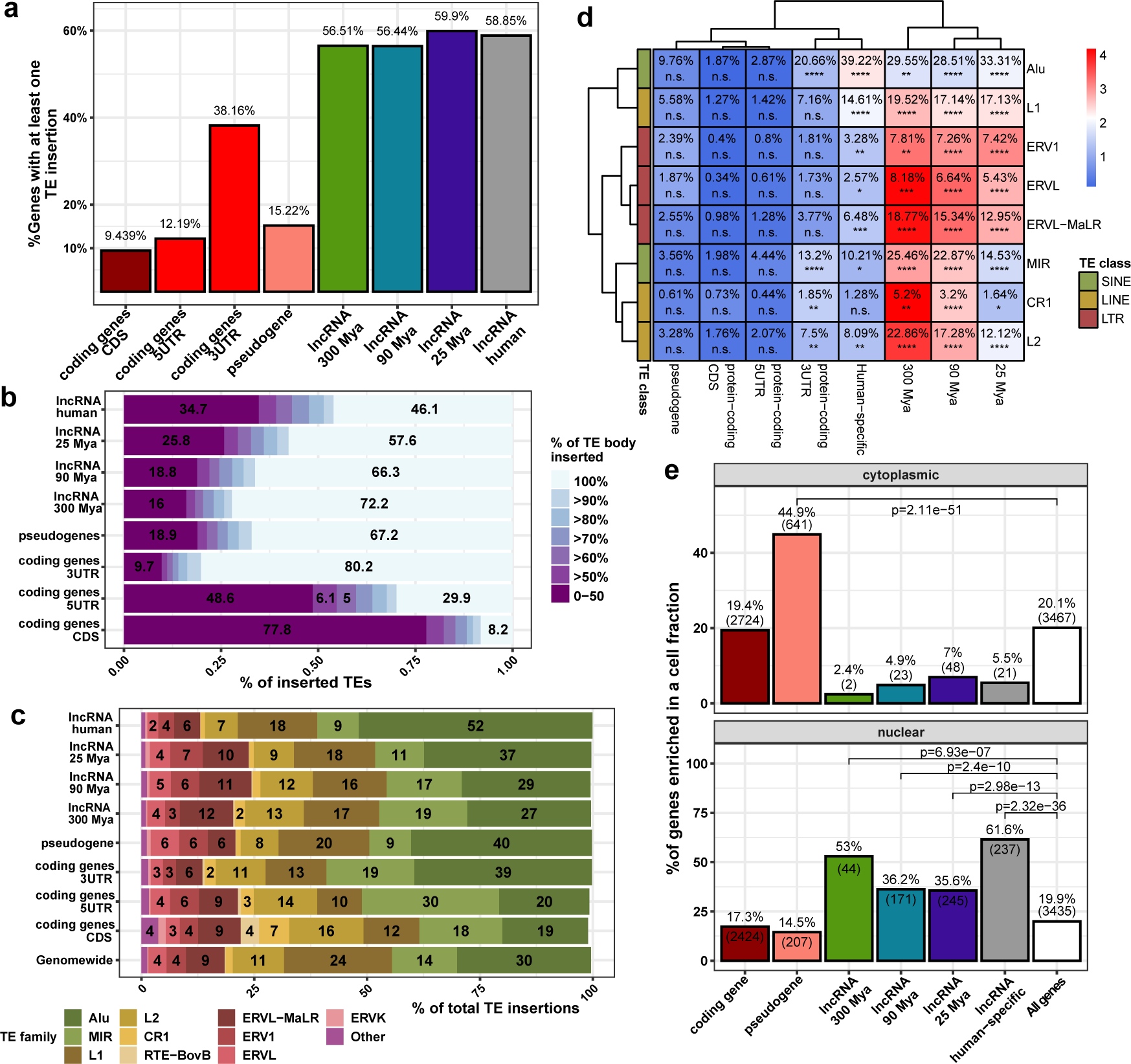
Transposable element insertion features of cortical lncRNAs throughout evolution. **a** Percentage of genes carrying at least one transposable element (TE) insertion. **b** For each gene type indicated at left, the horizontal bars show the fraction of inserted TEs with the indicated % of TE body inserted into the gene body (colored according to the scale at right). **c** Percentage of each TE family (color coded as shown at the bottom) inserted into each gene type indicated at left. **d** Heatmap displaying the percentage of genes in a category carrying at least one TE family insertion. Heatmap colors are proportional to the scaled percentage by each TE family. **e** Percentage of genes differentially expressed in the cellular compartment indicated at the top of each panel. P values are Fisher’s hypergeometric test. **Statistics**: All statistics labels are FDR-corrected P values of one-sided (greater) Fisher hypergeometric tests. *: p equal to or less than 0.05, **: p equal to or less than 10^-5^, ***: p equal to or less than 10^-10^, ****: p equal to or less than 10^-15^.

It has been hypothesized that exonic TEs have evolved as RNA domains essential for lncRNA function [30]. Thus, differences among lncRNA MA groups in the inserted TE family type might point to differences in their functionality. We assessed the distribution of TE families among the different gene categories to evaluate this possibility. All gene categories showed deviations from the overall genome-wide distribution of TE families (Fig. 3c). The LINE family L1, the second most abundant TE in the genome, was depleted from all considered gene types (Fig. 3c), especially from protein-coding genes; interestingly, pseudogenes are genes with the highest frequency of L1 insertion (Fig. 3c), pointing to L1 insertion as a mark of pseudogenization of coding genes. Moreover, most of the lncRNA categories, except the human-specific lncRNAs, showed a markedly higher frequency of endogenous retroviruses (ERVs), particularly the ERVL-MaLR family, which is the most abundant ERV in the human genome (Fig. 3c). Furthermore, we evaluated the percentage of genes carrying at least one TE family insertion among gene categories. We identified that ERVs are conspicuously inserted into lncRNAs, indicating that tolerance to ERV insertions is a feature of cortical lncRNAs (Fig. 3d).

Remarkably, L1 and Alu, active transposable elements of the human genome [31], are the TE families more frequently present in human-specific lncRNAs (Fig. 3c–d); in particular, Alu represents more than half of the TE insertion of this group of lncRNAs, while inactive TE families are enriched in older lncRNAs (Fig. 3c–d). This trend of TE family enrichment follows a gradient, where Alu occurrence among lncRNA MA groups is more prevalent in younger lncRNAs; inversely, inactive TE families such as ERVs, L2, CR1, and MIR are more prevalent in older lncRNAs than in younger lncRNAs (Fig. 3c–d).

Several TE families have been recognized as signals of nuclear retention of lncRNAs [32, 33]. Due to the differences in the distribution of TE families among the lncRNA MA groups, the nuclear retention feature of lncRNAs might also differ. Thus, we assessed the distribution of lncRNAs among the nuclear and cytoplasmic compartments in human fetal cortical tissues (Fig. 3e and Additional files 2 and 4: Tables S1 and S4). Remarkably, lncRNAs of all MA groups were proportionally more expressed in the nucleus than in the cytoplasm (Fig. 3e); regarding the lncRNAs expressed in the nucleus, different levels of expression resulted in lncRNA fractions varying from 35.6 to 61.6% among the different lncRNA MA groups. These results show that nuclear retention is a feature that has prevailed in lncRNAs through evolution.

### lncRNA evolutionary groups show distinct genomic distributions that highlight a potential functional specialization

The local genomic context of lncRNAs is intimately associated with their functionality [34, 35]. Thus, systematically inspecting protein-coding genes proximal to lncRNAs of different MA groups might shed light on the function that cortical lncRNAs have gained throughout evolution. For that, the nearest protein-coding genes were retrieved to a hundred kilobases surrounding the lncRNAs of different MA groups; then, we assessed their enriched GO terms. We found that MA groups evolved from loci near distinct types of developmental protein-coding genes that regulate different stages of neuron specification and maturation (Fig. 4a and Additional file 4: Table S5). Thus, ancient lncRNAs that appeared before the evolution of the cerebral cortex (300 Mya) are preferentially expressed from loci near broad developmental genes, including TFs (Fig. 4a, left and Additional file 4: Table S5), which suggests a pleiotropic function of those lncRNAs; meanwhile, a fraction of lncRNAs that appeared before the expansion of the primate cerebral cortex (90 Mya) are preferentially expressed from loci near protein-coding genes associated with the development of axons (Fig. 4a, center and Additional file 4: Table S5), and a fraction of human-specific lncRNAs are preferentially expressed from loci proximal to genes associated with dendrite development, where synapses are finely tuned in response to environmental cues (Fig. 4a, right and Additional file 4: Table S5).

**Fig. 4.**
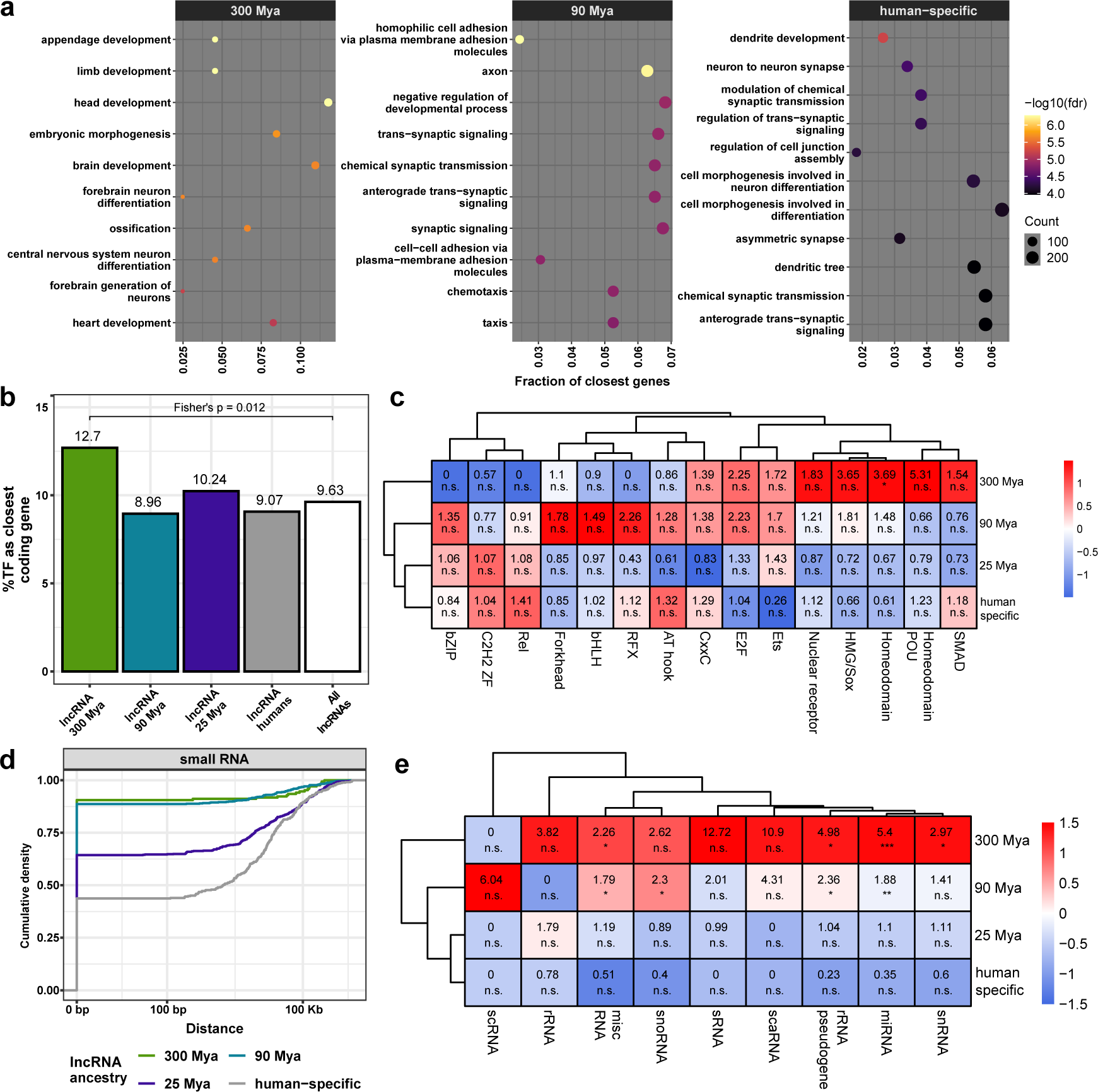
Differential genomic distribution of cortical lncRNAs. **a** Top 10 gene ontology terms enriched in the sets of protein-coding genes closest to the different lncRNA minimal-evolutionary-age (MA) groups indicated at the top of each panel. **b** Percentage of transcription factors (TF) as the closest genes for each lncRNA MA group. **c** Percentage of TF as the nearest gene separated by TF families, indicated in the columns; the displayed number indicates the odds ratio across of MA groups. Heatmap colors are proportional to the scaled odds ratio across TF family. **d** Cumulative distribution of the distance from the lncRNA to the nearest small RNA separated by MA groups. **e** Like **c** but for small RNAs as the closest gene. **Abbreviations:** snRNA, small nuclear RNA; miRNA, microRNA; rRNA, ribosomal RNA; scaRNA, small Cajal body-specific RNA; sRNA, small RNA; snoRNA, small nucleolar RNA; scRNA, small cytoplasmic RNA. **Statistics**: All statistics labels are FDR-corrected P values of one-sided (greater) Fisher hypergeometric tests. Ns: p greater than 0.05, *: p equal to or less than 0.05, **: p equal to or less than 10^-5^, ***: p equal to or less than 10^-10^.

Furthermore, one of the enriched GO terms of protein-coding genes from the vicinity of ancient cortical lncRNAs was “DNA-binding transcription activator activity” (Additional file 4: Table S5); therefore, we tested whether older lncRNAs have an enriched number of TFs in their proximity, finding that only the older MA group has a slightly increased abundance of proximal TFs compared with the other MA groups (Fig. 4b). Due to the considerable fraction of TFs as the closest coding gene (8.96 to 12.7%) and the fact that TFs are the master regulators of biological functioning, we sought to identify the type of TFs proximal to lncRNAs of different MA groups. Surprisingly, lncRNAs of different MA groups are preferentially distributed around certain families of TFs. Older lncRNAs are preferentially expressed from loci close to the homeodomain-containing TFs (Fig. 4c), master regulators of early development. Meanwhile, newer lncRNAs are preferentially expressed from loci near C2H2 zinc finger-containing (ZF) TFs, a fast-evolving family of transcriptional repressors [36, 37] (Fig. 4c).

We observed that older lncRNAs (300 and 90 Mya) are more often proximal to small RNAs than younger lncRNAs (25 Mya and human-specific) (Fig. 4d); it has been shown that several lncRNAs host small RNAs [38]. Therefore, we tested whether older lncRNAs were sources of small RNA genes and found that older lncRNAs preferentially host small RNAs in their loci (Fig. 4e), especially microRNAs, which indicates a possible coevolution of older cortical lncRNAs with microRNAs.

### lncRNA evolutionary groups exhibit different expression patterns and cellular enrichment

We have shown that cortical lncRNAs display commonalities, differentiating them from mRNAs and pseudogenes. They also show differences among MA groups (locus complexity, TE composition, and genomic distribution) that point to a shift in their functionality throughout evolution. However, the extent to which lncRNA evolution might impact the biology of the cerebral cortex remains unexplored.

To start evaluating the possible biological function of lncRNAs from distinct MA groups in the developing cerebral cortex, we assessed the expression pattern of lncRNAs from the different MA groups in a set of mid-gestational RNA-seq libraries (Additional files 2 and 4: Table S1 and S6). We identified that younger lncRNAs are significantly depleted from the cortical germinative zones but not from the cortical plate (Fig. 5a). To further explore the cellular context in which lncRNAs from different MA groups may function, we reanalyzed public scRNA-seq data from the developing human cerebral cortex [27, 28], mapping the cortical lncRNAs to the cellular populations of the human developing cerebral cortex. All identified cell populations (Fig. 5b) differentially expressed at least one member of each MA group (Fig. 5c Additional file 4: Table S6), indicating widespread expression of all cortical lncRNAs. Of note, lncRNAs are enriched in mature excitatory neurons (Cajal-Retzius, synaptogenic glutamatergic, and mature subplate neurons), and several cortical cell populations preferentially express lncRNAs from certain MA groups. Thus, ancient lncRNAs (300 Mya) are enriched in all interneuron cell populations (Inh. CGE, Inh. SST, Inh. MGE), outer RGCs, and migrating glutamatergic neurons; instead, young lncRNAs (25 Mya, human-specific) were preferentially enriched in mature subplate (SP) and synaptogenic glutamatergic neurons (Fig. 5c). In line with the depletion from germinative zones, cortical lncRNAs are depleted from cycling cell populations (apical and outer RGCs, mGPC, IPCs, microglia, oligodendrocytes, astrocytes, OPC), except for ancient lncRNAs that are enriched in oRGCs; at the same time, protein-coding genes are also enriched in those cell populations (Fig. 5c).

**Fig. 5.**
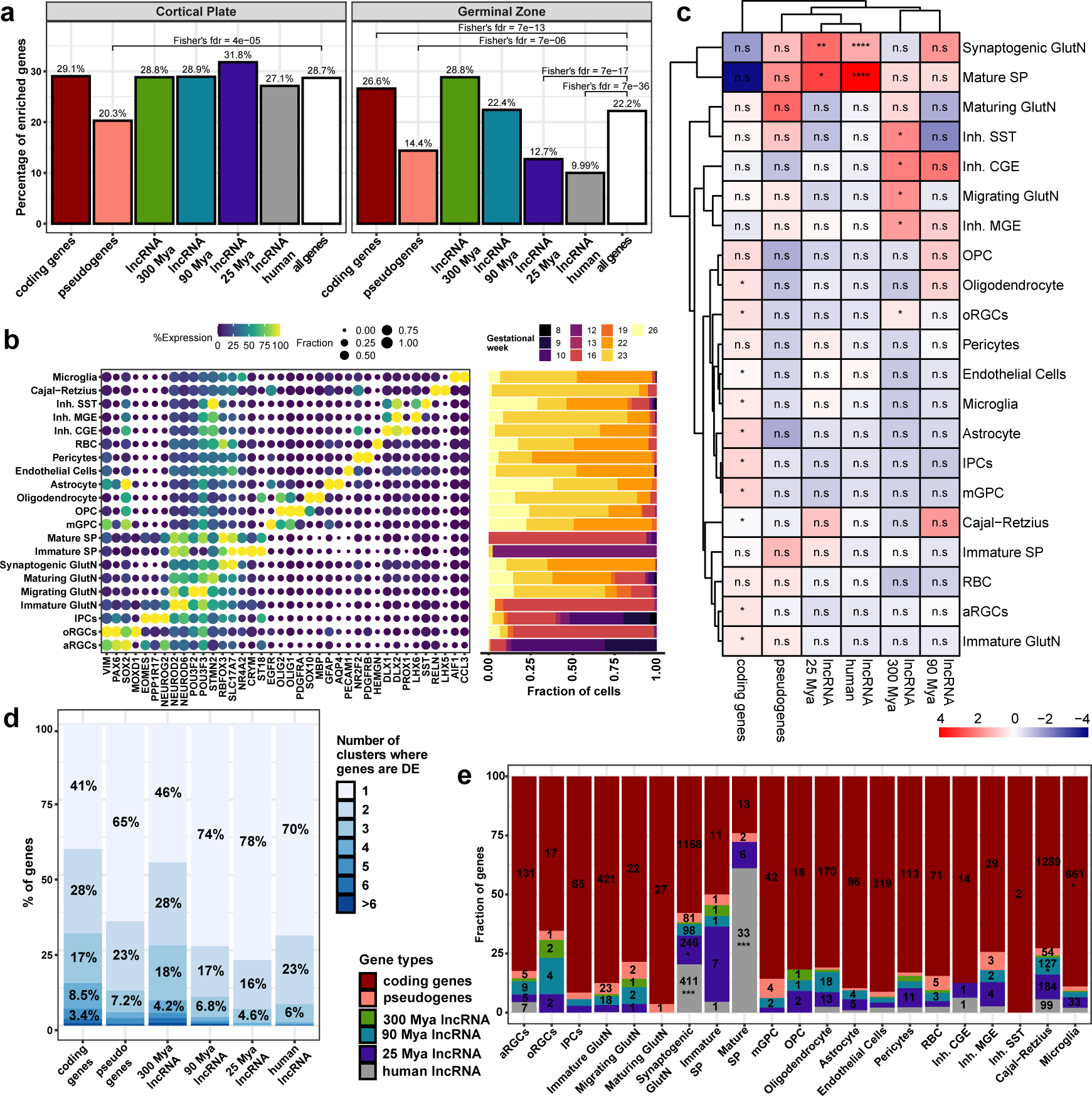
Differential distribution of lncRNAs in cortical cell types throughout evolution. **a** Percentage of differentially expressed (DE) enriched genes by gene category in cortical plate (left) or germinal (right) zones of mid-gestation cerebral cortices. FDR values on the enriched gene categories come from Fisher’s hypergeometric test. **b** Dotplot (left) showing the average expression of the cell population marker genes (x-axis) related to the single-cell cluster (y-axis) with the maximum expression, and barplot (right) displaying the fraction of cells from a cluster coming from a gestational week (right). **c** Heatmap displaying the MA groups (columns) identified in our analyses as enriched in each cell type (indicated on the right). Heatmap colors are proportional to the z-score across MA groups of the percentage of the distinct gene types in each cell type. **d** Frequency of the cell type-specificity of different gene types; for each gene type (column), the percentage of those genes that are DE in 1 to 6 or more cell clusters (as indicated by the color) is given inside the boxes; these DE genes are specific markers of those cell clusters. **e** Frequency of cell type-specific genes colored by gene type (color scale at left). Cell types are indicated at the x-axis; numbers inside the bars indicate the absolute number of cell type-specific lncRNAs in each cell type cluster. **Abbreviations:** aRGCs, apical radial glial cells; oRGCs, outer radial glial cells; IPCs, intermediate progenitor cells; GlutN, glutamatergic neurons; SP, subplate neurons; mGPC, multipotent glial progenitor cells; OPC, oligodendrocyte progenitor cells; RBC, red blood cells; Inh. CGE, inhibitory GABAergic interneurons derived from the caudal ganglionic eminences; Inh. MGE, inhibitory GABAergic interneurons derived from the medial ganglionic eminences; Inh. SST, inhibitory GABAergic interneurons expressing somatostatin. **Statistics**: All statistics labels are FDR-corrected P values of one-sided (greater) Fisher hypergeometric tests. ns: p greater than 0.05, *: p equal to or less than 0.05, **: p equal to or less than 10^-5^, ***: p equal to or less than 10^-10^, ****: p equal to or less than 10^-15^.

It has been shown that lncRNAs with low expression in bulk RNA-seq data are specifically expressed in a single cell population in the developing cerebral cortex [39]. Therefore, we assessed the specificity of cortical lncRNA expression in all gene categories. Of all gene categories considered, protein-coding genes were the most broadly expressed among cell clusters, as only 41% of protein-coding genes were specific to a single cell population (Fig. 5d). Among lncRNA MA groups, ancient lncRNAs were shown to be more broadly expressed among several cell populations (only 46% of lncRNAs from this MA group are specific to a single cell cluster). In comparison, younger lncRNAs share a similarly high percentage of cell-type specificity (78% – 70%) (Fig. 5d). Next, we assessed the cell-type enrichment of the cell-type-specific lncRNAs (Fig. 4e) and found that Cajal-Retzius cells, mature subplate (SP) neurons, and synaptogenic glutamatergic neurons were the only cell types expressing significantly higher absolute numbers of cell type-specific lncRNAs compared with other cells (Fig. 5e). Of the cell type-specific lncRNAs, large numbers of mammalian, primate, and human-specific lncRNAs were enriched in Cajal-Retzius cells, while primate (25 Mya) and human-specific lncRNAs were enriched in synaptogenic glutamatergic neurons; finally, mature SP neurons were enriched in human-specific lncRNAs (Fig. 5e).

### Cortical lncRNAs are regulated by a set of specific transcription factors

To better understand the molecular basis of the differential cellular distribution of cortical lncRNAs through evolution, we assessed their regulatory landscape using the Remap data [40]. We found that cortical lncRNAs present a higher number of transcription factors (TFs) bound to their promoters than a set of random genomic regions and pseudogenes (Fig. 6a), reducing the possibility that most of the identified lncRNAs were bioinformatic artifacts. Collectively, all cortical lncRNAs present a reduced diversity of TFs in their promoters compared to protein-coding genes. Nevertheless, older lncRNAs (300 Mya and 90 Mya) show a higher variety of TFs than younger lncRNAs (Fig. 6a), indicating that older lncRNAs are under more complex regulation.

**Fig. 6.**
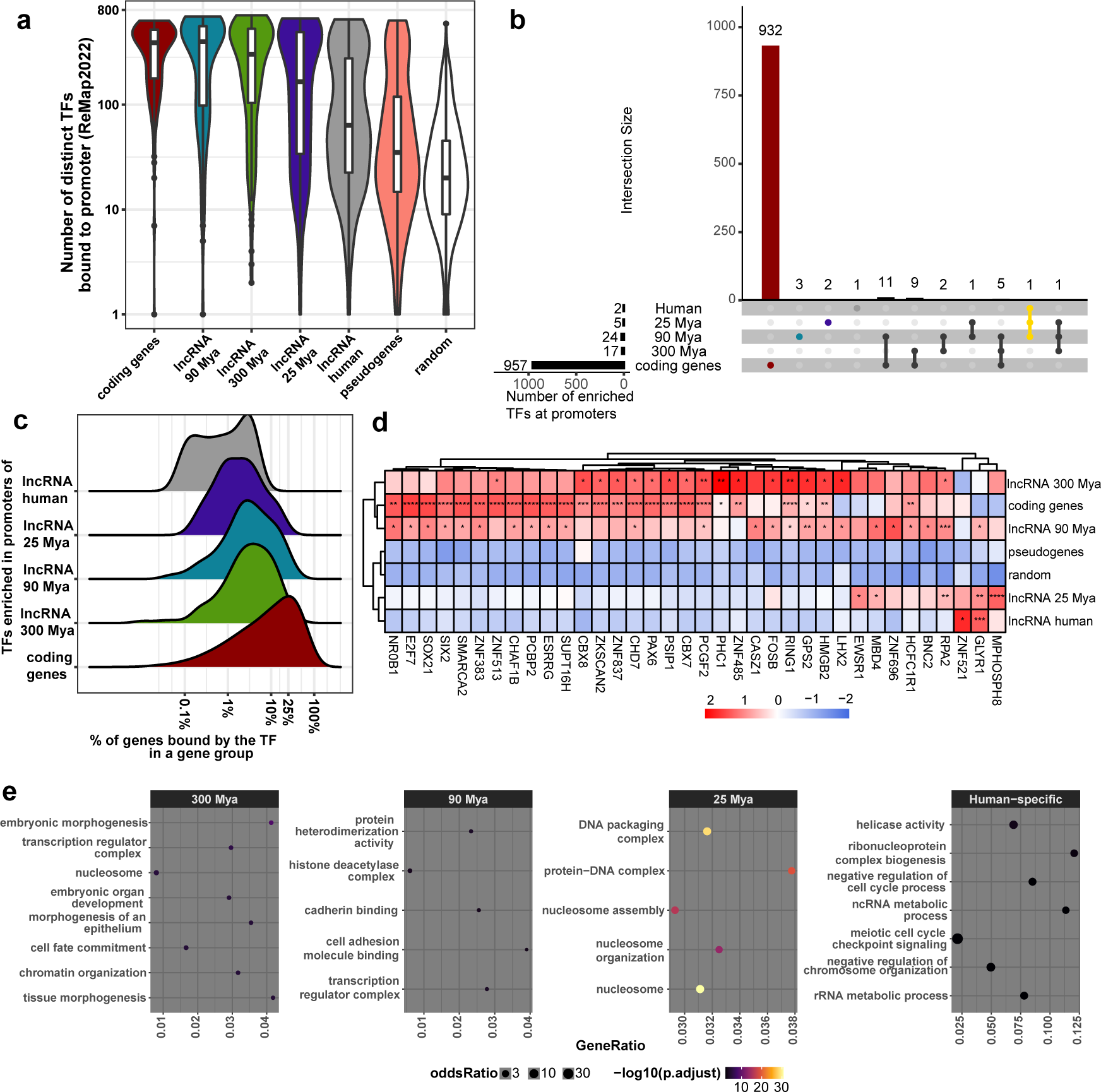
Regulatory landscape of cortical lncRNAs. **a** Number of different transcription factors (TFs) bound to the gene promoter region of genes in the gene category indicated at the x-axis. **b** UpSet plot displaying the number of shared and specific TFs enriched in promoters of lncRNA minimal-evolutionary-age (MA) groups and of protein-coding genes. **c** Distribution of the frequency of genes in all gene groups having an enriched TF bound in their promoter. **d** Heatmap of the frequency of TFs bound in promoters of gene groups. Heatmap colors are proportional to the scaled frequency by gene group. **e** Gene ontology enrichment of protein-coding genes having TFs enriched in the promoter of different lncRNA MA groups bound to their promoters. **Statistics**: All statistics labels are FDR-corrected P values of one-sided (greater) Fisher hypergeometric tests*: p equal to or less than 0.05, **: p equal to or less than 10^-5^, ***: p equal to or less than 10^-10^, ****: p equal to or less than 10^-15^.

Next, we looked for TFs enriched in promoters of lncRNA MA groups. We found that most TFs are preferentially bound to promoters of protein-coding genes, few of them in promoters of lncRNAs, many of which were enriched in promoters of 300 and 90 Mya MA groups and shared with TFs in promoters of protein-coding genes (Fig. 6b), in concordance with the higher diversity of TFs found in older lncRNA promoters. Interestingly, TFs enriched in all lncRNA MA groups regulate fewer numbers of genes in both protein-coding and lncRNA MA groups, indicating that lncRNA regulators, in general, are more specific (Fig. 6c). Furthermore, we found that TFs enriched in young lncRNAs (25 Mya and human-specific) were depleted from protein-coding gene promoters (Fig. 6d), implying that these TFs preferentially regulate lncRNAs. Additionally, we noticed that protein-coding genes regulated by TFs also present in common at the promoters of lncRNAs from each MA group are part of different molecular pathways (Fig. 6e). TFs present in promoters of antique lncRNAs (300 Mya) preferentially regulate protein-coding genes associated with embryonic morphogenesis, TFs present in promoters of mammalian-specific lncRNAs (90 Mya) preferentially regulate genes associated with histone deacetylase and cadherin binding, and TFs present in promoters of primate-specific lncRNAs (25 Mya) preferentially regulate histone genes (Fig. 6e). Finally, TFs present in promoters of human-specific lncRNAs preferentially control genes that negatively regulate the cell cycle process (Fig. 6e and Additional file 5: Table S7).

Due to their specificity for regulating lncRNAs, we explored the set of TFs enriched in promoters of young lncRNAs. Most of the TFs enriched in promoters of primate-specific lncRNAs, namely, EWSR1, MDB4, and RPA2, regulated protein-coding genes involved in the nucleosome assembly process (Additional file 1: Fig. S5a and Additional file 5: Table S8). Two of them, EWSR1 and RPA2, are expressed in intermediate progenitor cells (IPCs, Additional file 1: Fig. S5b), indicating that these TFs are involved in the regulation of proliferation, and due to the depletion of primate-specific lncRNAs from this cell type, they might negatively regulate the expression of primate-specific lncRNAs in IPCs. Furthermore, the TF GLYR1 is progressively enriched in mammalian, primate, and human-specific lncRNA promoters and is upregulated in synaptogenic glutamatergic neurons and Cajal-Retzius cells (Fig. 6d and Additional file 1: Fig. S5b). As GLYR1 is a known transcriptional activator, it may regulate the overexpression of lncRNAs in mature neurons. Moreover, GLYR1 preferentially regulated genes involved in the negative regulation of the cell cycle (Additional file 1: Fig. S5a and Additional file 5: Table S8). The sharing of positive regulation of genes that stop or slow cell proliferation together with regulation of lncRNAs may explain the reduced expression of lncRNAs in proliferative cell populations and their increased number in nonmitotic mature neurons.

### Young lncRNAs are sources of molecular innovation of cortical excitatory neurons

To understand how lncRNAs have impacted the gene coexpression networks of human corticogenesis, we employed weighted gene coexpression network analysis (WGCNA) [41]. After filtering library outliers, we used 187 bulk RNA-seq samples from the cerebral cortex that expanded from prenatal to early postconception, from which we identified 43 gene coexpression modules (Additional file 5: Table S9). Then, we assessed the centrality of the different lncRNA MA groups in these modules by comparing their intramodular connectivity (kIM) distribution. Protein-coding genes were significantly more central than all lncRNA MA groups (Fig. 7a). At the same time, cortical lncRNAs followed a gradient, where older lncRNAs were more central than younger lncRNAs, with human-specific lncRNAs following a distribution similar to pseudogenes (Fig. 7a). These results indicate that protein-coding genes are pivotal for maintaining coexpression networks. At the same time, lncRNA MA groups usually build up around central protein-coding genes following an ontological pattern, tuning the expression modules.

**Fig. 7.**
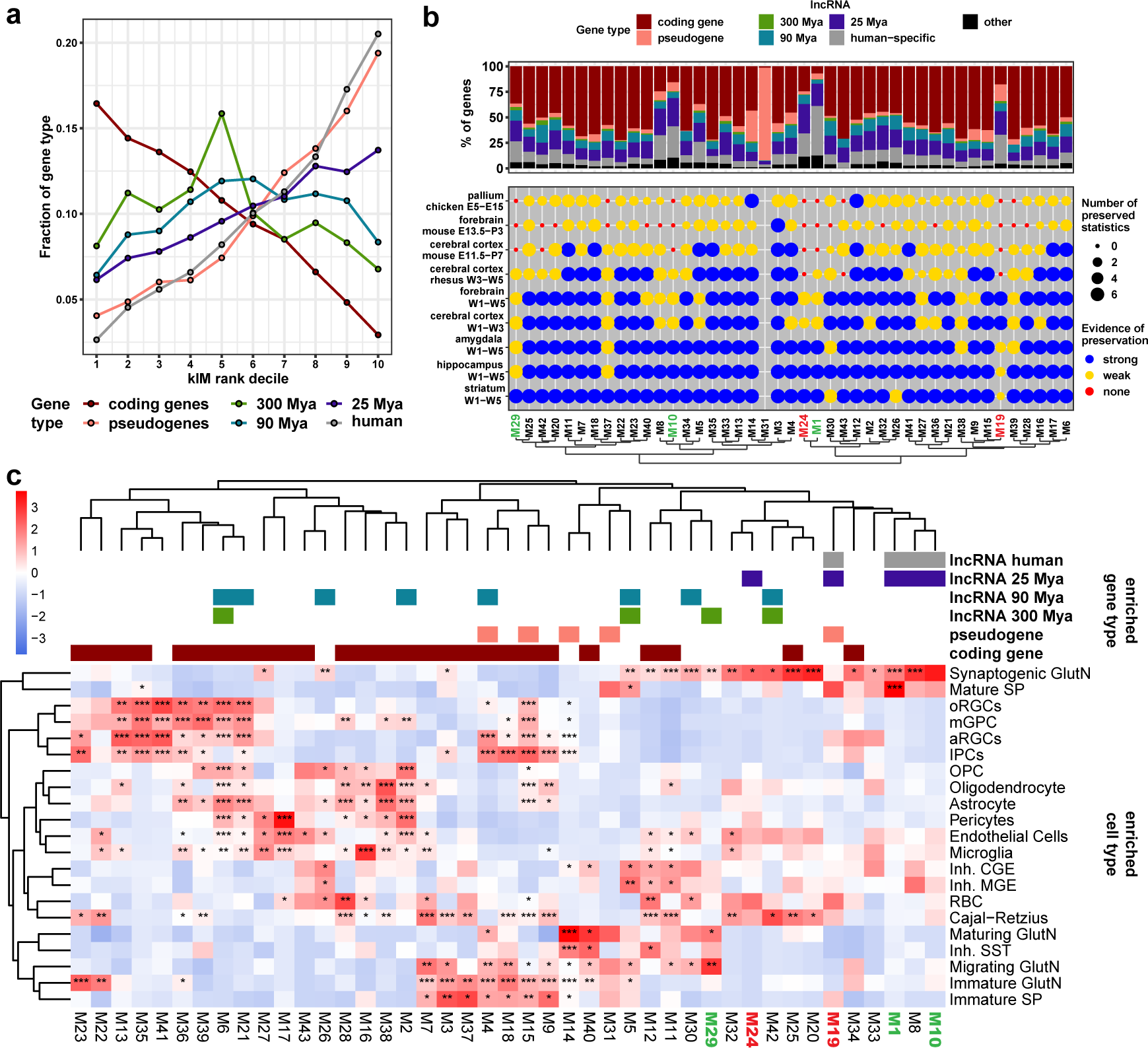
Young cortical lncRNAs contribute to novel molecular features of the developing cerebral cortex. **a** Fraction of genes (y-axis) from each gene category (colored as indicated at the bottom) present at the given intramodular connectivity (kIM) decile (x-axis). In weighted gene coexpression modules, kIM indicates how well a gene is connected to other members of the gene modules. The first kIM corresponds to genes with the highest connectivity to other genes from the same gene coexpression module. **b** Preservation analysis of 7 module statistics of cortical modules in coexpression networks from human, macaque, mouse, and chicken forebrain tissues. **Strongly** preserved modules: all 7 module statistics were found preserved (Bonferroni adjusted P value < 0.05); **weak**: between 6 and 1 module statistics were found preserved; **none**: no module feature was found preserved. Fractions of gene types are displayed in the top panel. **c** Intersection of cortical modules and scRNA-seq DE data, where the heatmap displays the odds ratio scaled by row. Colored blocks at the top indicate that a lncRNA minimal-evolutionary-age group (MA) indicated at the top right is enriched in that gene module. Asterisk labels correspond to the statistical significance of the gene module enrichment at the given cortical cell types indicated on the right. **Statistics**: All statistics labels are FDR-corrected P values of one-sided (greater) Fisher hypergeometric tests. *: p equal to or less than 0.05, **: p equal to or less than 10^-5^, ***: p equal to or less than 10^-10^, ****: p equal to or less than 10^-15^.

Furthermore, we assessed the role of cortical lncRNAs in gene network innovation using preservation network analyses, comparing seven network properties [42] of the human developing cortex gene coexpression modules with the coexpression networks of developing tissues of the human, macaque, mouse, and chicken forebrains (Additional file 5: Table S10). We found that modules M19 and M24 were preserved only in humans and not in any other of the assessed species (Fig. 7b, red dots in chicken pallium, mouse forebrain and cortex, and rhesus cortex); as well, modules M1, M10, and M29 were only weakly preserved with the rhesus macaque (Fig. 7b, yellow dots only in rhesus cortex); therefore, we identified modules M19 and M24 as human specific, while modules M1, M10, and M29 were only primate specific. Remarkably, most primate-and human-specific gene modules, namely, M1, M10, M19, and M24, were also enriched in younger lncRNAs (Fig. 7b, top blue and grey color bars). Furthermore, when intersecting with the scRNA-seq DE data, it was possible to identify the cortical cell types enriched in these more divergent modules; in particular, excitatory neurons (synaptogenic glutamatergic and mature SP neurons) are highly enriched in primate and human-specific coexpression modules (Fig. 7c), thus indicating that *de novo* expression of lncRNAs has preferentially been involved in the molecular diversification of excitatory neurons.

### Human-specific lncRNAs are dysregulated in autism spectrum disorder (ASD)

Several studies have shown that neuropsychiatric disorders disrupt the homeostasis of postmitotic developing glutamatergic neurons [18, 43, 44]. As young lncRNAs are conspicuously expressed in this cell population and involved in its molecular diversification, we investigated whether cortical lncRNAs are dysregulated in neuropsychiatric disorders. Public bulk RNA-seq data from cortical tissues of normal and autism spectrum disorder (ASD) specimens [45] were subjected to DE analysis (Fig. 8a and Additional file 5: Table S11). Remarkably, we identified that young cortical (primate and human-specific) lncRNAs are highly dysregulated in ASD (Fig. 8a and 8b), and these upregulated young lncRNAs are preferentially expressed in synaptogenic glutamatergic and mature subplate neurons (Fig. 8c). Despite the high dysregulation of young cortical lncRNAs, especially human-specific lncRNAs, we did not identify differences in the expression of the most proximal protein-coding genes (Fig. 8d), suggesting lncRNAs possibly acting in trans. Nevertheless, proximal protein-coding genes to dysregulated young lncRNAs are involved in neurite development and synapse formation (Fig. 8e Additional file 5: Table S12), two disrupted functions in ASD.

**Fig. 8.**
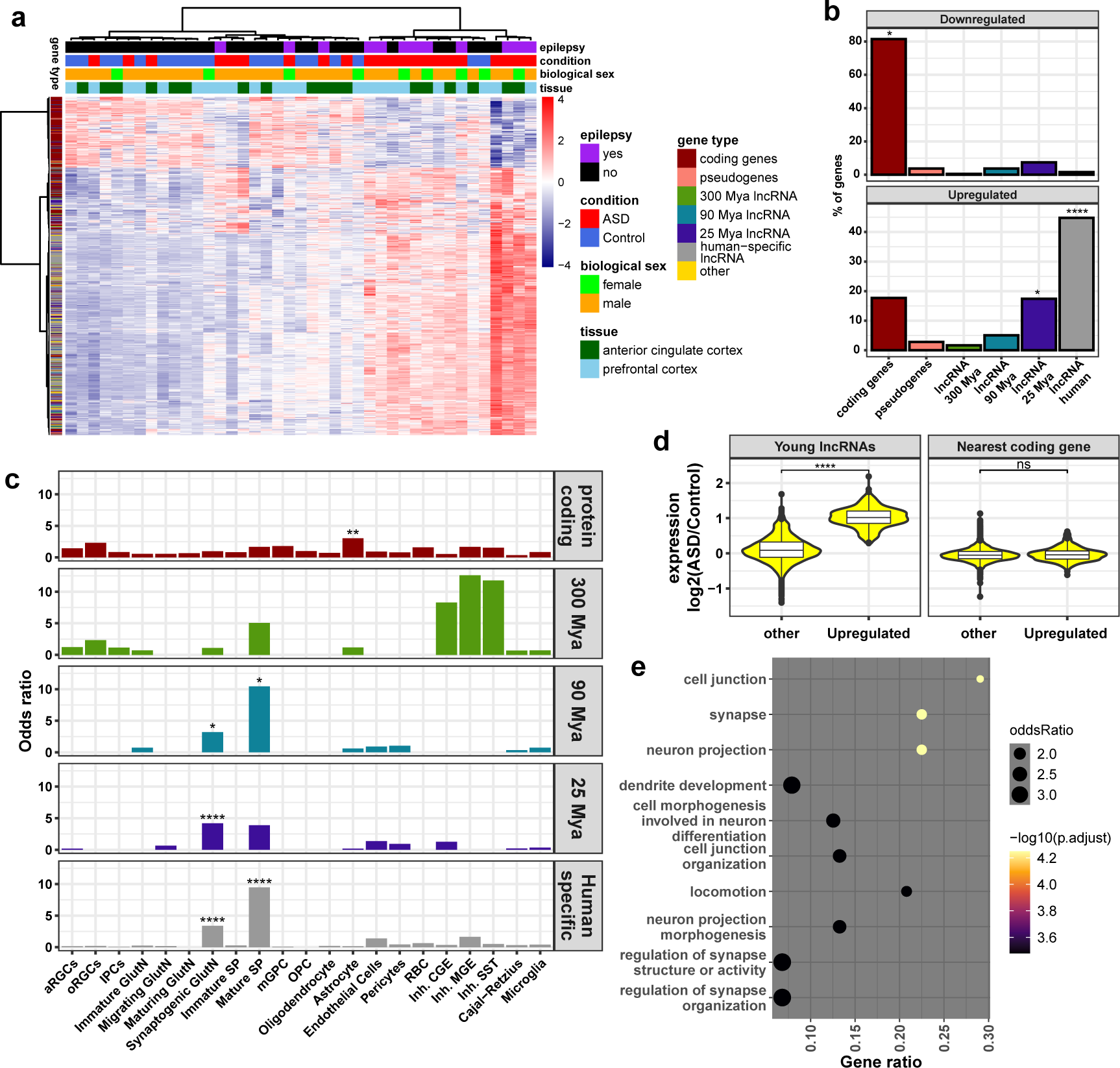
Young lncRNAs are dysregulated in ASD. **a** Heatmap (unsupervised clustering) of differentially expressed genes in samples from autism spectrum disorders (ASD) compared with control individuals. The heatmap color indicates the z-score for each gene (line) calculated across all samples (columns) and is indicated by the color scale on the right. Sample characteristics are indicated by colors in the top panels and specified in the legend on the right. Gene types are shown in the far-left column, and their color legend is on the right. **b** Percentage of differentially expressed genes, either downregulated (upper panel) or upregulated (lower panel) in ASD relative to control samples, belonging to a gene category: protein-coding genes, pseudogenes, or one of lncRNA minimal-evolutionary-age (MA) groups. **c** Enrichment of upregulated genes in specimens with ASD expressed in the cell types of the developing cerebral cortex indicated in the x-axis, separated by gene category (each of the five panels). **d** Log2-fold change expression differences of upregulated young lncRNAs (primate and human-specific, left panel) in specimens with ASD and other (nondifferentially expressed) genes, and expression of their most proximal protein-coding genes (right panel). **e** Gene ontology enrichment of most proximal protein-coding genes of young cortical lncRNAs upregulated in specimens with ASD. **Statistics**: Labels represent FDR-corrected Fisher hypergeometric P values. ns: p greater than 0.05, *: p equal to or less than 0.05, **: p equal to or less than 10^-5^, ***: p equal to or less than 10^-10^, ****: p equal to or less than 10^-15^.

## Discussion

### Comprehensive annotation of lncRNAs

The low expression and tissue-specific features of lncRNAs make them challenging to annotate. Additionally, corticogenesis is a highly dynamic process that integrates many cells from different developmental regions; in particular, human corticogenesis is lengthy and comprises more diverse cells than other mammals [1, 2]. Consequently, many lowly expressed lncRNAs or lncRNAs restricted to rare cell types of the developing cerebral cortex might not be annotated in public references such as Gencode [46] or Ensembl [24], which is detrimental to studying the evolution of lncRNAs and limits our understanding of the transcriptional complexity of this tissue. A critical contribution of the present work was annotating a comprehensive complement of lncRNAs for humans, rhesus macaques, mice, and chickens. The large number of libraries comprising the different periods of prenatal brain development in the four studied species (Additional file 1: Fig. S1) enabled us to identify new lncRNAs, protein-coding genes, and pseudogenes and to increase the number of reads mapped to the annotated transcriptomes compared to reference annotations (Additional file 1: Fig. S4 i–l), representing a qualitative and quantitative improvement.

### Estimation of lncRNA minimal age

To characterize the evolution of cortical lncRNA features, we estimated the minimal evolutionary age for all human lncRNAs based on their syntenic conservation across amniotes. The concept of syntenic conservation of lncRNAs as an indicator of homology [10, 25] was combined with the reciprocal conservation of genetic elements to avoid the misidentification of syntenic lncRNAs. Due to the stringent criteria, we incorporated syntenic conservation annotation of lncRNAs from previous studies [10, 12, 26] to help reduce the possibility of misclassifying lncRNAs into evolutionary groups. We documented the accuracy of the MA groups by examining different features. As expected, we found that older lncRNAs present higher evolutionary conservation scores than younger lncRNAs (Fig. 1e). Additionally, older lncRNAs display higher expression levels and increased splicing efficiency and have longer transcripts than younger lncRNAs (Fig. 2b, 2f, and 2g). These are features previously associated with conserved lncRNAs [10, 12, 47]. Furthermore, we documented a distinct distribution of TEs in lncRNAs, with younger lncRNAs having a progressively higher frequency of primate-specific Alu insertions (Fig. 3c and 3d), which has been linked with the formation of primate neurological networks [48]. Finally, we showed a depletion of younger lncRNAs in germinative zones and their increased expression in postmitotic neurons. This aligns with the reported high expression of species-specific lncRNAs at late prenatal stages [12]. In summary, these features support the accuracy of the MA classification and the sequential ages they represent.

### Cortical lncRNAs display features of functional specialization

Due to their reduced evolutionary constraints and expression levels, the functionality of lncRNAs has been questioned. Previous systematic analyses of lncRNAs in vertebrates have shown that these RNAs present features of positive selection, including higher conservation than random sequences, expression regulation by TFs, splicing activity, and dynamic expression [11, 12, 49]. We showed that our cohort of cortical lncRNAs presents similar signatures of positive selection, namely, having higher evolutionary constraints than intronic protein-coding regions (except for human-specific lncRNAs), canonic splicing, a higher number of distinct TFs bound at their promoters than pseudogenes, and cell type-specific expression (Fig. 1e, 2f, 6a, 5e, respectively). Nevertheless, these signatures differ among the MA groups, with young lncRNAs displaying weaker selection signals than older lncRNAs (Fig. 2b–h and 6a). In addition, the low expression levels of lncRNAs are associated with their steady degradation by exosome complexes, which leads to their accumulation in chromatin [50]. We found that the differences in expression levels among MA groups were notorious, with antique lncRNAs having as much expression as mRNAs (Fig. 2b); in addition, all MA groups were enriched in the nucleus (Fig. 3e). Furthermore, older lncRNAs present longer transcripts, shorter exons, larger introns, and more isoforms (Fig. 2g, 2d, 2e, and 2h, respectively), which indicate selection against early transcriptional termination and alternative splicing regulation. Overall, these distinct features among MA groups point to increased functionality of the RNA product of older lncRNAs; meanwhile, the shorter, younger lncRNAs might not primarily act in an RNA-dependent manner. These differences are consistent with the proposed constructive neutral evolution regime of lncRNAs, which postulated that *de novo* expression of lncRNAs by pervasive transcription of RNA pol II or TE transposition is first maintained by the selection of RNA transcription *per se* and then by additional modularity gain during evolution [51]. That said, we understand that not necessarily all lncRNAs identified in this study have a measurable phenotype; nevertheless, it is more likely that older lncRNAs have gained one.

In addition to the different functional capabilities among MA groups, we showed that multiple genomic features point to a shift in the role of cortical lncRNAs throughout evolution. We found that antique lncRNAs are regulated by TFs that control protein-coding genes involved in broad developmental processes (Fig. 6e). Additionally, antique lncRNAs are not depleted from cortical germinative zones and have a wider expression in different cell types of the developing cerebral cortex than younger lncRNAs (Fig. 5a, 5c–d). Moreover, antique lncRNAs exhibit high insertion of ERVs, are typically expressed in embryonic and fetal development [52] (Fig. 3c–d) and are proximal to developmental regulatory genes such as homeobox TFs and microRNAs (Fig. 4). These results indicate that antique lncRNAs have evolved along with and likely act as regulators of conserved developmental programs through various mechanisms, including serving as a source of or competing with microRNA or acting as *trans* and *cis* regulators [7]. A good example of an antique cortical lncRNA is *DLX6-AS1*, a conserved lncRNA with two functionally characterized isoforms that regulate the development of inhibitory neurons by controlling the expression of nearby *DLX5/6* TFs [53]. Remarkably, we found that antique lncRNAs are enriched in inhibitory cortical neurons (Fig. 5c), a conserved neuron type of the cerebral cortex [54]. Another cell type enriched in antique lncRNAs is the outer radial glial cells (oRGCs, Fig. 5c), a neural progenitor type substantially expanded in human corticogenesis [1]. Then, antique lncRNAs may coevolve with preserved developmental programs to assist in diversifying conserved cell types.

On the other hand, cortical lncRNAs that evolved after the divergence of amniotes are preferentially expressed near genes that regulate neurite development, are depleted from germinative zones, and are conspicuously expressed in excitatory neurons (Cajal-Retzius cells, Sub Plate cells, and glutamatergic neurons) (Fig. 4a, 5a, 5c, and 5e, respectively). Moreover, primate- and human-specific lncRNAs are regulated by a few TFs that showed a higher frequency in lncRNAs and coregulated protein-coding genes negatively associated with cell cycle progression, which explains their increased cell-type selectivity for excitatory neurons and their depletion from germinative zones. Furthermore, we found that young cortical lncRNAs integrate human-specific gene coexpression modules enriched in mature glutamatergic neurons (Fig. 7b–c). These features indicate that these lncRNAs are more relevant for the proper development and molecular diversification of mature excitatory neurons. It is tempting to postulate that young unstable lncRNAs, which are enriched in the proximity of genes or in introns of genes (Fig. 2a) involved in axon and dendrite development (Fig. 4a), may interact with template DNA of RNA pol II to form R-loops to relax chromatin and allow prolonged transcription. In line with this idea, it was recently found that Alu sequences from eRNAs and upstream antisense lncRNAs help to mediate enhancer-promoter interactions [55]. This could be an interesting mechanism of action for young lncRNAs, as the human cerebral cortex is characterized by protracted development compared to other primates and mammals in general [1]. Then, young cortical lncRNAs could be used by excitatory neurons to sustain prolonged transcription throughout their extended development. Furthermore, the increased expression of young lncRNAs in ASD, a human-specific disease characterized by defective synapsis, indicates that they may be important for the proper synaptogenesis of human neurons. Of note, we did not identify differences in the bulk expression levels of protein-coding genes nearest to lncRNAs affected in ASD, which does not eliminate a possible function of these young lncRNAs in mRNA transcription and processing, modulation of specific protein‒protein or protein‒ligand interactions, or differences at the single-cell RNA level. Thus, we postulate that young cortical lncRNAs have been used as a molecular source of evolution and are involved in the disease of human excitatory neurons, primarily in synaptogenesis.

## Conclusions

In this study, we systematically assessed the contribution of lncRNAs to human cerebral cortex evolution by annotating comprehensive catalogs of cortical lncRNAs in humans, rhesus macaques, mice, and chickens and by identifying the minimal evolutionary age of human cortical lncRNAs. We showed that human cortical lncRNAs exhibit signatures of functional specialization throughout evolution. Older lncRNAs are preferentially expressed near developmental regulatory genes such as homeobox TFs and microRNAs and are enriched in conserved cell types, such as oRGCs and inhibitory neurons, working as a source of molecular innovation of conserved molecular programs. On the other hand, younger cortical lncRNAs are selectively regulated to be expressed in postmitotic excitatory neurons, divergent neurons of the cerebral cortex, where they form primate and human-specific coexpression networks and are dysregulated in autism spectrum disorder, indicating that they are a source of molecular evolution and dysfunction of excitatory neurons.

## Acknowledgments

We thank Dr. C. Y. Irene Yan, Instituto de Ciências Biomédicas, Universidade de São Paulo, for expert advice and discussions on aspects of developmental biology. Data used here, which had been generated as part of the PsychENCODE Consortium, was supported by grants: U01DA048279, U01MH103339, U01MH103340, U01MH103346, U01MH103365, U01MH103392, U01MH116438, U01MH116441, U01MH116442, U01MH116488, U01MH116489, U01MH116492, U01MH122590, U01MH122591, U01MH122592, U01MH122849, U01MH122678, U01MH122681, U01MH116487, U01MH122509, R01MH094714, R01MH105472, R01MH105898, R01MH109677, R01MH109715, R01MH110905, R01MH110920, R01MH110921, R01MH110926, R01MH110927, R01MH110928, R01MH111721, R01MH117291, R01MH117292, R01MH117293, R21MH102791, R21MH103877, R21MH105853, R21MH105881, R21MH109956, R56MH114899, R56MH114901, R56MH114911, R01MH125516, and P50MH106934 awarded to: Alexej Abyzov, Nadav Ahituv, Schahram Akbarian, Alexander Arguello, Lora Bingaman, Kristin Brennand, Andrew Chess, Gregory Cooper, Gregory Crawford, Stella Dracheva, Peggy Farnham, Mark Gerstein, Daniel Geschwind, Fernando Goes, Vahram Haroutunian, Thomas M. Hyde, Andrew Jaffe, Peng Jin, Manolis Kellis, Joel Kleinman, James A. Knowles, Arnold Kriegstein, Chunyu Liu, Keri Martinowich, Eran Mukamel, Richard Myers, Charles Nemeroff, Mette Peters, Dalila Pinto, Katherine Pollard, Kerry Ressler, Panos Roussos, Stephan Sanders, Nenad Sestan, Pamela Sklar, Nick Sokol, Matthew State, Jason Stein, Patrick Sullivan, Flora Vaccarino, Stephen Warren, Daniel Weinberger, Sherman Weissman, Zhiping Weng, Kevin White, A. Jeremy Willsey, Hyejung Won, and Peter Zandi.

## Authors’ contributions

Conceptualization: DAMV, SVA. Methodology: DAMV. Formal analysis: DAMV. Investigation: DAMV, ACT, MSA. Resources: DAMV, DWS, MSA, MGBC, SVA. Data Curation: DAM. Writing original draft: DAMV. Writing review and editing: DAMV, ACT, DWS, MSA, MGBC, SVA. Visualization: DAMV. Funding acquisition: SVA. Project administration: SVA. Supervision: ACT, SVA.

## Data availability

Raw chicken RNA-seq data sequenced for this project were deposited at the NCBI SRA repository under Accession Number PRJNA1002381. The list of public library Accession Numbers from different repositories (<u>Synapse</u> and <u>SRA</u>) used in the present work can be found in Additional file 2: Table S1. The new transcriptome annotations, the identified minimal evolutionary age of human lncRNAs, and their matching homologous lncRNAs in the other studied species are provided in Additional file 3: Table S2. GTF files with the transcriptome annotations for all species used in the present project can be found at https://doi.org/10.5281/zenodo.10038371.

## Methods

### Tissue collection, RNA extraction, and sequencing

*Gallus gallus* fertilized eggs were purchased from a local provider and incubated at 38 °C and 50% humidity for seven and ten days. Embryos were collected and decapitated; brains were removed from the heads, and forebrains were further dissected. For developmental day 7 (E7), the whole pallium and subpallium were retrieved; for developmental day 10 (E10), the entire subpallium, the dorsolateral pallium, and the medial pallium were retrieved. Three brain sections from different embryos were pooled per working sample without considering biological sex. Brain sections were dissociated using pestles in 1 ml TRIzol and frozen at -80 °C until the day of RNA isolation.

For RNA isolation, 200 µL of chloroform was added per 1 ml TRIzol and centrifuged for 15 min and 16000 g at 4 °C; supernatants were transferred to new microtubes. One volume of 100% ethanol was added to each sample and then transferred to RNeasy Mini spin columns. After that, the RNeasy Microkit protocol was followed.

RNA samples were quantified using a Qubit2 Fluorometer (Thermo Fisher), and their integrity was measured using a Bioanalyzer 2100 (Agilent). The RNA integrity number (RIN) of the samples ranged from 7.5 to 8.5, indicating good quality. Four biological replicates were prepared and sent for sequencing to BGI Genomics (Shenzhen, China) for each tissue-developmental window.

### Bulk RNA-seq processing

Public libraries were retrieved from the SRA repository at GenBank (NCBI, USA) using *fasterq-dump* with the following parameter: “--split-files”; the integrity of the data was checked using *vdb-validate*, and all files were identified as consistent. Mapped bam files for the rhesus macaques were retrieved from the Synapse repository (https://www.synapse.org/#!Synapse:syn17093056/tables/RhesusmRNA-seq) using the repository API for UNIX and then transformed into fastq files using the *bedtools bamtofastq* [56] function. Raw fastq files generated in the present work and sequenced at BGI Genomics were retrieved from a dedicated AMAZON Web services account. Raw fastq files from all sources were then processed with *fastp* [57] to remove read adapters and to check read quality before and after trimming. Trimmed fastq files were mapped to the reference genome using *STAR* version 2.5.4b [58] using the following parameters: “--outReadsUnmapped Fastx --chimSegmentMin 12 -- chimJunctionOverhangMin 12 --alignSJDBoverhangMin 10 –alignMatesGapMax 100000 --alignIntronMax 100000 --chimSegmentReadGapMax 3 -- alignSJstitchMismatchNmax 5 -1 5 5 --runThreadN 94 --outSAMstrandField intronMotif --outFilterMultimapNmax 20 --outFilterType BySJout -- outFilterMismatchNoverReadLmax 0.04 --alignIntronMin 20 --outSAMtype BAM Unsorted”. The latest available genomes were used as genome references (hg38, rheMac10 [20], mm39 [21], and galGal6 [22, 23]). The resulting unsorted BAM files were sorted using *samtools* [59].

### Iso-seq long-read processing

Raw unmapped bam files from the SRA project PRJNA476474 were downloaded directly from the SRA repository using GNU wget for the rhesus macaque. The standard *Isoseq3* pipeline was used to obtain polished, high-quality fasta files for all processed samples. Additionally, for the chicken (*Gallus gallus*), fasta files from Iso-seq sequencing deposited at SRA were downloaded using *fasterq-dump*, as described above. Detailed information on all libraries used can be found in Additional file 2: Table S1.

Long reads fasta files were mapped to the reference genomes using *Minimap2* [60] with the following parameters: “-ax splice -uf --secondary=no -C5 -O6,24 -B4”. Output sam files were converted to bam files, sorted, and indexed using *samtools*. All output sam files from the same species were collapsed into a gtf file using the function *collapse_isoforms_by_sam.py* from *Cupcake* with default parameters. Spurious transcripts were removed from the collapsed gtf file using the functions *sqanti3_qc.py and sqanti3_RulesFilter.py* from *Squnti3* [61] with default parameters.

### GTF building

To generate consensus gene models from short reads, sorted bam files from bulk RNA-seq libraries were processed using *scallop* [62] with the following parameters: “--min_transcript_length_base 200 --min_mapping_quality 250 --min_splice_bundary_hits 1”. To choose the correct parameter for “--library_type”, the type of library was assessed before running *scallop*; for the parameter “--min_num_hits_in_bundle”, 10 was chosen if the library possessed less than 20 million uniquely mapped reads; otherwise, 20 was used. After generating gtf files for every sample, the monoexonic transcripts were removed from unstranded libraries. Additionally, the monoexonic transcripts were removed from rRNA-depleted libraries using *gffread* [63] with the parameter “gffread in_gtf_file -F -U -T -o /out_put/file”.

GTF files from all samples of the same developmental window/brain region were merged into a consensus transcriptome using *taco* [64] before generating the final gtf files to avoid overrepresentation of libraries of a tissue/brain region group, which would bias the construction of the final transcript model, with the following parameters “--gtf-expr-attr RPKM --filter-min-expr 0 --filter-min-length 200 --isoform-frac 0.01”. Consensus transcriptomes were merged into a final gtf file using *taco* with similar parameters. The output consensus file was filtered for readthrough, mapping errors, intron retention, and run-on polymerase transcripts using *gffcompare* [63] with the species reference transcriptome as a model to generate the final consensus gtf file.

### Coding potential identification

Coding potential was assessed for all transcripts in the final gtf files using *CPAT3* [65], *FEELnc* [66], and *CPC2* [67]. For CPAT and CPC2, gtf files were transformed into fasta files, first generating intermediate bed files with *gffread*, then using *getfasta* from *samtools* with the following parameters: “-split -name -s.” The fasta files were used as input to generate coding potential values for each transcript. For FEELnc, the reference gtf files for each evaluated species were used to train the random forest model, running the tool with the following parameters: “-n 6000,6000 --learnorftype=3 --testorftype=3”. Tables with output coding potential scores can be found in Additional file 3: Table S2.

### Open reading frame identification and annotation

*Transdecoder* 5.5.0 [68] was used to identify *bona fide* ORFs; additional Pfam and UniProt matches were provided to improve the identification of ORFs. The *HMMER* 3.1b2 tool [69] was used to identify Pfam matches with the following parameters: “hmmscan --domtblout pfam.domtblout --tblout file_name.tsv -E 1e-5”. To identify UniProt matches, *blastp* was used with the following parameters: “-max_target_seqs 1 - outfmt 6 -evalue 1e-5”. The final identification of ORFs was carried out using the function *TransDecoder. Predict* with the following parameters: “--retain_pfam_hits pfam.domtblout --retain_blastp_hits blastp.outfmt6”. Additionally, gff3 files were generated for each species containing the genomic coordinates of the ORFs, information that was added to the final consensus transcriptomes. Finally, eggNOG-mapper [70] matches were identified for all identified ORFs using the UNIX standalone tool *emapper* 2.0.1 with default parameters. Output annotations of identified ORFs are incorporated in Additional file 3: Table S2.

### Transposable element content identification

Repetitive elements from each studied species were downloaded from the UCSC Genome Browser database (https://genome.ucsc.edu), keeping only the records from transposable element (TE) families SINE, LINE, LTR, DNA, Retroposon, and RC. TE tables were converted to bed files using custom *Rscripts* and sorted using UNIX *sort*.

To identify ORFs with more than 50% of their gene body coming from a TE element, protein-coding sequence (CDS) bed files were intersected with TE bed files using the *bedtools intersect* function with the following parameters: “-s -wo” to ensure strand specificity of the intersection. The total sum of TE intersections was divided by the length of the CDS; CDSs with more than 50% of their gene body coming from a TE element were tagged as “Transposable elements.”

To identify the TE class distribution in CDS, mRNA untranslated regions (UTRs), pseudogenes, and lncRNAs, the bed files from the new assemblies of each species were intersected with the TE bed file. TEs that intersected at least ten bp with a gene were kept for further analysis. The percentage of the gene body containing TEs was calculated as the ratio of the total length of all TEs intersecting with a gene to the gene length.

### Identification of syntenic lncRNAs

Two approaches were used to identify syntenic conserved lncRNAs between humans and the other studied species. Whole genome alignment and long transcript mapping. In both cases, all isoforms from a gene were merged into a metagene annotation, generating new bed files of metagenes. Due to the repetitive nature of TEs, we removed them from metagene models using *bedtools subtract*. FASTA files for metagene annotations without TEs were generated using *bedtools getfasta*. In the whole genome alignment approach (*liftover*), metagene bed files were lifted to the genome coordinates of the other species using the standalone *liftover* function from the UCSC Genome Browser with the following parameters: “-minBlocks=0.01 -minMatch=0.01”. In the transcript mapping approach, metagene fasta files were mapped to the genome of the other species using *Minimap2* with the following parameters: “-ax splice -uf -- secondary=no -C5 -O4,24 -A2 -B4 -G 100K”. The output sam files were converted to bam sorted files using *samtools* and then to bed files using *bedtools bamtobed*.

Bed files containing mapped lncRNAs from one species to the other genome were joined, prioritizing the transferred genes using the whole genome alignment approach. The final set of transferred gene coordinates was intersected with the group of lncRNAs from the target species. LncRNAs with the same position and orientation and that had at least one neighboring one-to-one protein-coding ortholog in the same direction were classified as syntenic lncRNAs. TEs enormously contribute to the gene body of lncRNAs [14]; removing TEs from the gene body of lncRNAs might lead to the misidentification of some syntenic lncRNAs. Thus, the same approaches were undertaken without removing the TE insertions from the lncRNA body. The syntenic information result from this analysis was added to the final annotation when an older syntenic lncRNA was identified concerning the previous analysis. For further analysis, only reciprocal syntenic conserved lncRNAs were used. Homologous lncRNAs and syntenic classification of lncRNAs are incorporated in Additional file 3: Table S2.

### Bulk RNA-seq quantification and differential expression analysis

Gene expression was quantified by *FeatureCounts* from the *Rsubread* package [71] using the new assemblies for each assessed species as a reference with the following parameters: “allowMultiOverlap = T, countMultiMappingReads = F, juncCounts = T, nthreads = 96”. The raw expression matrix was batch-corrected for humans using ComBat-seq [72], as in the original PsychEncode publication [18]. TPM values were calculated from raw expression matrices as previously shown by [73]:

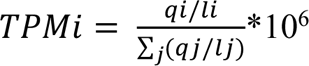

where *TPMi* is the TPM value of *gene i*, *qi* is the number of reads mapped in *gene i*, *li* is the length in kilobases of *gene i*, and ∑_j_(*qj/lj*) is the sum of counts/length ratios of all genes.

Only genes with a TPM value greater than 0.5 for all samples from developmental window/brain region pairs were kept for the subsequent analyses. Filtered TPM matrices were normalized using variance stabilization normalization [74].

To identify differentially expressed genes (DEGs), the R package edgeR [75] was used. Briefly, lowly expressed genes (less than 0.5 CPM in all samples of a variable) from raw count matrices were removed. Filtered matrices were used to identify DEGs using the quasilikelihood test; the resulting P values were FDR corrected, and all genes with an FDR less than 0.05 were identified as DEGs.

### Single-cell RNA-seq processing

Fastq files were retrieved from SRA using fasterq-dump as described above but with the following parameters: “fasterq-dump -S -O /output/dir -e 94 --include-technical”. Fastq files were then mapped to the reference genome using *STARsolo* [76] version 2.7.9a with the following parameters: “--soloType CB_UMI_Simple –soloCBwhitelist /barcodes/dir --soloBarcodeMate 0 --soloBarcodeReadLength 0 --soloCBstart 1 -- soloCBlen 8 --soloUMIstart 9 --soloUMIlen 8 --readFilesCommand zcat --runThreadN 94 --soloStrand Forward --clipAdapterType CellRanger4 --readFilesIn READ1.fq READ2.fq --soloCellFilter None --soloFeatures Gene Velocyto GeneFull -- soloMultiMappers PropUnique”. Raw, sparse matrices from all samples were loaded into R [77] and merged using *Seurat* [78]. Cells with fewer than ten thousand and more than one million UMIs, fewer than a thousand detected genes, and more than 5% of all counts mapped to mitochondrial genes were removed from further analysis. Raw, sparse expression matrix was normalized using SCT transformation while regressing by the percentage of expressed mitochondrial genes, cellular-cycle score, number of UMIs, and the number of identified genes using the following parameters: “method = “glmGamPoi,” vst.flavor = “v2”, variable.features.n = 5000, vars.to.regress = c(“percent.mt”, “CC.Difference”, “nCount_RNA”, “nFeature_RNA”).”

Single-cell clusters were identified using the Seurat function FindNeighbors considering the first fifty dimensions and the function FindClusters with resolution *2.5*. Markers for all clusters were identified using the functions FindAllMarkers with the following parameters: “assay = “SCT,” test.use = “wilcox”, only.pos = T, logfc.threshold = 0.25, min.pct = .25, return.thresh = 0.05, densify = T”. Known cell population gene markers were used to annotate the identified clusters to different cortical cell types.

### Identification of expression-matched genes

The R package “optmatch” [79] was used to identify the set of expression-matched genes among the evaluated gene categories, utilizing the mean-variance stabilized expression of all samples from the same gene as input.

### Identification of genes closest to lncRNAs

The *bedtools* function *closest* was used to identify the most proximal genes with the following parameters: “-d,” for which a metagene bed file for protein-coding genes and small RNAs was built for each species. Only genes located in the genome approximately 100 kb of lncRNAs were kept for the following analysis. Gene Ontology (GO) analysis of the closest protein-coding genes was undertaken using the R package clusterProfiler [80, 81] and using as input the list of nearest protein-coding genes to a minimal age (MA) category and the list of all closest protein-coding genes as background.

### Transcription factor assay

Promoters from the most extensive transcript with the highest number of exons were retrieved from a set of expression-matched genes from all assessed gene categories. An equal number of random genomic regions were also generated using *bedtools shuffle* as a null group. Nonredundant remap 2022 data [40] were retrieved and intersected with the set of working promoters using *bedtools intersect*. The absolute number of different proteins bound to each gene category and random genomic regions were compared using the Wilcoxon test. To identify enriched TFs in the promoter of a gene category, a one-sided Fisher hypergeometric test was performed between the set of random sequences and the promoters from the gene category. P values were FDR corrected, and all TFs with an FDR less than 0.05 were identified as enriched in that gene category. To test the function of the TFs enriched in each gene category, all genes with the TF bound to their promoter were retrieved and used for GO analysis, using all genes with at least one TF enriched in their promoter as background.

### Statistical Analysis

All statistical plots and tests were obtained using the statistical package R version 4.1.0 [77].

## References

1. Libé-Philippot B, Vanderhaeghen P. Cellular and Molecular Mechanisms Linking Human Cortical Development and Evolution. Annu Rev Genet 2021;55:555–81.

2. Molnár Z, Clowry GJ, Šestan N, Alzu’bi A, Bakken T, Hevner RF, Hüppi PS, Kostović I, Rakic P, Anton ES, et al. New insights into the development of the human cerebral cortex. J Anat 2019;235(3):432–51.

3. Silbereis John C, Pochareddy S, Zhu Y, Li M, Sestan N. The Cellular and Molecular Landscapes of the Developing Human Central Nervous System. Neuron 2016;89(2):248–68.

4. Lui JH, Hansen DV, Kriegstein AR. Development and evolution of the human neocortex. Cell 2011;146(1):18–36.

5. Berg J, Sorensen SA, Ting JT, Miller JA, Chartrand T, Buchin A, Bakken TE, Budzillo A, Dee N, Ding S-L, et al. Human neocortical expansion involves glutamatergic neuron diversification. Nature 2021;598(7879):151–8.

6. Vanderhaeghen P, Polleux F. Developmental mechanisms underlying the evolution of human cortical circuits. Nature Reviews Neuroscience 2023;24(4):213–32.

7. Statello L, Guo C-J, Chen L-L, Huarte M. Gene regulation by long non-coding RNAs and its biological functions. Nature Reviews Molecular Cell Biology 2021;22(2):96–118.

8. Ransohoff JD, Wei Y, Khavari PA. The functions and unique features of long intergenic non-coding RNA. Nat Rev Mol Cell Biol 2018;19(3):143–57.

9. Rinn JL, Chang HY. Long Noncoding RNAs: Molecular Modalities to Organismal Functions. Annu Rev Biochem 2020;89:283–308.

10. Hezroni H, Koppstein D, Schwartz MG, Avrutin A, Bartel DP, Ulitsky I. Principles of long noncoding RNA evolution derived from direct comparison of transcriptomes in 17 species. Cell reports 2015;11(7):1110–22.

11. Necsulea A, Soumillon M, Warnefors M, Liechti A, Daish T, Zeller U, Baker JC, Grutzner F, Kaessmann H. The evolution of lncRNA repertoires and expression patterns in tetrapods. Nature 2014;505(7485):635–40.

12. Sarropoulos I, Marin R, Cardoso-Moreira M, Kaessmann H. Developmental dynamics of lncRNAs across mammalian organs and species. Nature 2019;571(7766):510–4.

13. Aprea J, Calegari F. Long non-coding RNAs in corticogenesis: deciphering the non-coding code of the brain. Embo j 2015;34(23):2865–84.

14. Kapusta A, Kronenberg Z, Lynch VJ, Zhuo X, Ramsay L, Bourque G, Yandell M, Feschotte C. Transposable Elements Are Major Contributors to the Origin, Diversification, and Regulation of Vertebrate Long Noncoding RNAs. PLOS Genetics 2013;9(4):e1003470.

15. Pollard KS, Salama SR, Lambert N, Lambot M-A, Coppens S, Pedersen JS, Katzman S, King B, Onodera C, Siepel A, et al. An RNA gene expressed during cortical development evolved rapidly in humans. Nature 2006;443(7108):167–72.

16. Guo CJ, Ma XK, Xing YH, Zheng CC, Xu YF, Shan L, Zhang J, Wang S, Wang Y, Carmichael GG, et al. Distinct Processing of lncRNAs Contributes to Non-conserved Functions in Stem Cells. Cell 2020;181(3):621–36.e22.

17. Ruiz-Orera J, Messeguer X, Subirana JA, Alba MM. Long non-coding RNAs as a source of new peptides. Elife 2014;3:e03523.

18. Li M, Santpere G, Imamura Kawasawa Y, Evgrafov OV, Gulden FO, Pochareddy S, Sunkin SM, Li Z, Shin Y, Zhu Y, et al. Integrative functional genomic analysis of human brain development and neuropsychiatric risks. Science 2018;362(6420):eaat7615.

19. Zhu Y, Sousa AMM, Gao T, Skarica M, Li M, Santpere G, Esteller-Cucala P, Juan D, Ferrández-Peral L, Gulden FO, et al. Spatiotemporal transcriptomic divergence across human and macaque brain development. Science 2018;362(6420):eaat8077.

20. Warren WC, Harris RA, Haukness M, Fiddes IT, Murali SC, Fernandes J, Dishuck PC, Storer JM, Raveendran M, Hillier LW, et al. Sequence diversity analyses of an improved rhesus macaque genome enhance its biomedical utility. Science 2020;370(6523):eabc6617.

21. Church DM, Schneider VA, Graves T, Auger K, Cunningham F, Bouk N, Chen HC, Agarwala R, McLaren WM, Ritchie GR, et al. Modernizing reference genome assemblies. PLoS Biol 2011;9(7):e1001091.

22. Bellott DW, Skaletsky H, Cho TJ, Brown L, Locke D, Chen N, Galkina S, Pyntikova T, Koutseva N, Graves T, et al. Avian W and mammalian Y chromosomes convergently retained dosage-sensitive regulators. Nat Genet 2017;49(3):387–94.

23. Steffen M, Petti A, Aach J, D’Haeseleer P, Church G. Automated modelling of signal transduction networks. BMC Bioinformatics 2002;3:34.

24. Yates AD, Achuthan P, Akanni W, Allen J, Allen J, Alvarez-Jarreta J, Amode MR, Armean IM, Azov AG, Bennett R, et al. Ensembl 2020. Nucleic Acids Research 2019;48(D1):D682–D8.

25. Ulitsky I, Shkumatava A, Jan CH, Sive H, Bartel DP. Conserved function of lincRNAs in vertebrate embryonic development despite rapid sequence evolution. Cell 2011;147(7):1537–50.

26. Bryzghalov O, Szcześniak MW, Makałowska I. SyntDB: defining orthologues of human long noncoding RNAs across primates. Nucleic Acids Research 2020;48(D1):D238–D45.

27. Fan X, Dong J, Zhong S, Wei Y, Wu Q, Yan L, Yong J, Sun L, Wang X, Zhao Y, et al. Spatial transcriptomic survey of human embryonic cerebral cortex by single-cell RNA-seq analysis. Cell Research 2018;28(7):730–45.

28. Zhong S, Zhang S, Fan X, Wu Q, Yan L, Dong J, Zhang H, Li L, Sun L, Pan N, et al. A single-cell RNA-seq survey of the developmental landscape of the human prefrontal cortex. Nature 2018;555(7697):524-8.

29. Siepel A, Bejerano G, Pedersen JS, Hinrichs AS, Hou M, Rosenbloom K, Clawson H, Spieth J, Hillier LW, Richards S, et al. Evolutionarily conserved elements in vertebrate, insect, worm, and yeast genomes. Genome Research 2005;15(8):1034–50.

30. Johnson R, Guigó R. The RIDL hypothesis: transposable elements as functional domains of long noncoding RNAs. Rna 2014;20(7):959–76.

31. Mills RE, Bennett EA, Iskow RC, Devine SE. Which transposable elements are active in the human genome? Trends Genet 2007;23(4):183–91.

32. Carlevaro-Fita J, Polidori T, Das M, Navarro C, Zoller TI, Johnson R. Ancient exapted transposable elements promote nuclear enrichment of human long noncoding RNAs. Genome Res 2019;29(2):208–22.

33. Lubelsky Y, Ulitsky I. Sequences enriched in Alu repeats drive nuclear localization of long RNAs in human cells. Nature 2018;555(7694):107–11.

34. Amaral PP, Leonardi T, Han N, Viré E, Gascoigne DK, Arias-Carrasco R, Büscher M, Pandolfini L, Zhang A, Pluchino S, et al. Genomic positional conservation identifies topological anchor point RNAs linked to developmental loci. Genome Biology 2018;19(1):32.

35. Engreitz JM, Haines JE, Perez EM, Munson G, Chen J, Kane M, McDonel PE, Guttman M, Lander ES. Local regulation of gene expression by lncRNA promoters, transcription and splicing. Nature 2016;539(7629):452–5.

36. Tadepally HD, Burger G, Aubry M. Evolution of C2H2-zinc finger genes and subfamilies in mammals: species-specific duplication and loss of clusters, genes and effector domains. BMC Evol Biol 2008;8:176.

37. Najafabadi HS, Mnaimneh S, Schmitges FW, Garton M, Lam KN, Yang A, Albu M, Weirauch MT, Radovani E, Kim PM, et al. C2H2 zinc finger proteins greatly expand the human regulatory lexicon. Nat Biotechnol 2015;33(5):555–62.

38. Sun Q, Song YJ, Prasanth KV. One locus with two roles: microRNA-independent functions of microRNA-host-gene locus-encoded long noncoding RNAs. Wiley Interdiscip Rev RNA 2021;12(3):e1625.

39. Liu SJ, Nowakowski TJ, Pollen AA, Lui JH, Horlbeck MA, Attenello FJ, He D, Weissman JS, Kriegstein AR, Diaz AA, Lim DA. Single-cell analysis of long non-coding RNAs in the developing human neocortex. Genome Biology 2016;17(1):67.

40. Hammal F, de Langen P, Bergon A, Lopez F, Ballester B. ReMap 2022: a database of Human, Mouse, Drosophila and Arabidopsis regulatory regions from an integrative analysis of DNA-binding sequencing experiments. Nucleic Acids Research 2022;50(D1):D316–D25.

41. Langfelder P, Horvath S. WGCNA: an R package for weighted correlation network analysis. BMC Bioinformatics 2008;9(1):559.

42. Ritchie SC, Watts S, Fearnley LG, Holt KE, Abraham G, Inouye M. A Scalable Permutation Approach Reveals Replication and Preservation Patterns of Network Modules in Large Datasets. Cell Syst 2016;3(1):71–82.

43. Parikshak NN, Swarup V, Belgard TG, Irimia M, Ramaswami G, Gandal MJ, Hartl C, Leppa V, Ubieta LT, Huang J, et al. Genome-wide changes in lncRNA, splicing, and regional gene expression patterns in autism. Nature 2016;540(7633):423–7.

44. Ziffra RS, Kim CN, Ross JM, Wilfert A, Turner TN, Haeussler M, Casella AM, Przytycki PF, Keough KC, Shin D, et al. Single-cell epigenomics reveals mechanisms of human cortical development. Nature 2021;598(7879):205–13.

45. Velmeshev D, Schirmer L, Jung D, Haeussler M, Perez Y, Mayer S, Bhaduri A, Goyal N, Rowitch DH, Kriegstein AR. Single-cell genomics identifies cell type-specific molecular changes in autism. Science 2019;364(6441):685–9.

46. Frankish A, Diekhans M, Ferreira A-M, Johnson R, Jungreis I, Loveland J, Mudge JM, Sisu C, Wright J, Armstrong J, et al. GENCODE reference annotation for the human and mouse genomes. Nucleic Acids Research 2018;47(D1):D766–D73.

47. Necsulea A, Soumillon M, Warnefors M, Liechti A, Daish T, Zeller U, Baker JC, Grutzner F, Kaessmann H. The evolution of lncRNA repertoires and expression patterns in tetrapods. Nature 2014;505(7485):635–40.

48. Larsen PA, Hunnicutt KE, Larsen RJ, Yoder AD, Saunders AM. Warning SINEs: Alu elements, evolution of the human brain, and the spectrum of neurological disease. Chromosome Res 2018;26(1-2):93–111.

49. Ponjavic J, Ponting CP, Lunter G. Functionality or transcriptional noise? Evidence for selection within long noncoding RNAs. Genome Res 2007;17(5):556–65.

50. Schlackow M, Nojima T, Gomes T, Dhir A, Carmo-Fonseca M, Proudfoot NJ. Distinctive Patterns of Transcription and RNA Processing for Human lincRNAs. Mol Cell 2017;65(1):25–38.

51. Palazzo AF, Koonin EV. Functional Long Non-coding RNAs Evolve from Junk Transcripts. Cell 2020;183(5):1151–61.

52. Bakoulis S, Krautz R, Alcaraz N, Salvatore M, Andersson R. Endogenous retroviruses co-opted as divergently transcribed regulatory elements shape the regulatory landscape of embryonic stem cells. Nucleic Acids Research 2022;50(4):2111–27.

53. Cajigas I, Chakraborty A, Swyter KR, Luo H, Bastidas M, Nigro M, Morris ER, Chen S, VanGompel MJW, Leib D, et al. The Evf2 Ultraconserved Enhancer lncRNA Functionally and Spatially Organizes Megabase Distant Genes in the Developing Forebrain. Mol Cell 2018;71(6):956–72.e9.

54. Tosches MA. From Cell Types to an Integrated Understanding of Brain Evolution: The Case of the Cerebral Cortex. Annu Rev Cell Dev Biol 2021;37(1):495–517.

55. Liang L, Cao C, Ji L, Cai Z, Wang D, Ye R, Chen J, Yu X, Zhou J, Bai Z, et al. Complementary Alu sequences mediate enhancer-promoter selectivity. Nature 2023;619(7971):868–75.

56. Quinlan AR, Hall IM. BEDTools: a flexible suite of utilities for comparing genomic features. Bioinformatics 2010;26(6):841–2.

57. Chen S, Zhou Y, Chen Y, Gu J. fastp: an ultra-fast all-in-one FASTQ preprocessor. Bioinformatics 2018;34(17):i884–i90.

58. Dobin A, Davis CA, Schlesinger F, Drenkow J, Zaleski C, Jha S, Batut P, Chaisson M, Gingeras TR. STAR: ultrafast universal RNA-seq aligner. Bioinformatics 2013;29(1):15–21.

59. Li H, Handsaker B, Wysoker A, Fennell T, Ruan J, Homer N, Marth G, Abecasis G, Durbin R, Genome Project Data Processing S. The Sequence Alignment/Map format and SAMtools. Bioinformatics 2009;25(16):2078–9.

60. Li H. Minimap2: pairwise alignment for nucleotide sequences. Bioinformatics 2018;34(18):3094–100.

61. Tardaguila M, de la Fuente L, Marti C, Pereira C, Pardo-Palacios FJ, Del Risco H, Ferrell M, Mellado M, Macchietto M, Verheggen K, et al. SQANTI: extensive characterization of long-read transcript sequences for quality control in full-length transcriptome identification and quantification. Genome Res 2018;28(3):396–411.

62. Shao M, Kingsford C. Accurate assembly of transcripts through phase-preserving graph decomposition. Nature Biotechnology 2017;35:1167.

63. Pertea G, Pertea M. GFF Utilities: GffRead and GffCompare. F1000Res 2020;9:304.

64. Niknafs YS, Pandian B, Iyer HK, Chinnaiyan AM, Iyer MK. TACO produces robust multisample transcriptome assemblies from RNA-seq. Nat Methods 2017;14(1):68–70.

65. Wang L, Park HJ, Dasari S, Wang S, Kocher JP, Li W. CPAT: Coding-Potential Assessment Tool using an alignment-free logistic regression model. Nucleic Acids Res 2013;41(6):e74.

66. Wucher V, Legeai F, Hedan B, Rizk G, Lagoutte L, Leeb T, Jagannathan V, Cadieu E, David A, Lohi H, et al. FEELnc: a tool for long non-coding RNA annotation and its application to the dog transcriptome. Nucleic Acids Res 2017;45(8):e57.

67. Kang YJ, Yang DC, Kong L, Hou M, Meng YQ, Wei L, Gao G. CPC2: a fast and accurate coding potential calculator based on sequence intrinsic features. Nucleic Acids Res 2017;45(W1):W12–W6.

68. Douglas P. TransDecoder https://github.com/TransDecoder/TransDecoder/wiki. 2018.

69. Mistry J, Finn RD, Eddy SR, Bateman A, Punta M. Challenges in homology search: HMMER3 and convergent evolution of coiled-coil regions. Nucleic Acids Res 2013;41(12):e121.

70. Huerta-Cepas J, Forslund K, Coelho LP, Szklarczyk D, Jensen LJ, von Mering C, Bork P. Fast Genome-Wide Functional Annotation through Orthology Assignment by eggNOG-Mapper. Molecular Biology and Evolution 2017;34(8):2115–22.

71. Liao Y, Smyth GK, Shi W. The R package Rsubread is easier, faster, cheaper and better for alignment and quantification of RNA sequencing reads. Nucleic Acids Res 2019;47(8):e47.

72. Zhang Y, Parmigiani G, Johnson WE. ComBat-seq: batch effect adjustment for RNA-seq count data. NAR Genomics and Bioinformatics 2020;2(3):lqaa078.

73. Zhao Y, Li M-C, Konaté MM, Chen L, Das B, Karlovich C, Williams PM, Evrard YA, Doroshow JH, McShane LM. TPM, FPKM, or Normalized Counts? A Comparative Study of Quantification Measures for the Analysis of RNA-seq Data from the NCI Patient-Derived Models Repository. Journal of Translational Medicine 2021;19(1):269.

74. Huber W, von Heydebreck A, Sültmann H, Poustka A, Vingron M. Variance stabilization applied to microarray data calibration and to the quantification of differential expression. Bioinformatics 2002;18(suppl_1):S96–S104.

75. McCarthy DJ, Chen Y, Smyth GK. Differential expression analysis of multifactor RNA-Seq experiments with respect to biological variation. Nucleic Acids Research 2012;40(10):4288–97.

76. Kaminow B, Yunusov D, Dobin A. STARsolo: accurate, fast and versatile mapping/quantification of single-cell and single-nucleus RNA-seq data. bioRxiv 2021(10.1101/2021.05.05.442755).

77. R: A Language and Environment for Statistical Computing [https://www.R-project.org/]

78. Hafemeister C, Satija R. Normalization and variance stabilization of single-cell RNA-seq data using regularized negative binomial regression. Genome Biology 2019;20(1):296.

79. Hansen BB, Klopfer SO. Optimal Full Matching and Related Designs via Network Flows. Journal of Computational and Graphical Statistics 2006;15(3):609–27.

80. Yu G, Wang L-G, Han Y, He Q-Y. clusterProfiler: an R Package for Comparing Biological Themes Among Gene Clusters. OMICS: A Journal of Integrative Biology 2012;16(5):284–7.

81. Wu T, Hu E, Xu S, Chen M, Guo P, Dai Z, Feng T, Zhou L, Tang W, Zhan L, et al. clusterProfiler 4.0: A universal enrichment tool for interpreting omics data. Innovation (Camb) 2021;2(3):100141.

