## Additional file 1_Suppl Figs S1 to S5 for "The human developing cerebral cortex is characterized by an increased de novo expression of lncRNAs in excitatory neurons"

### Additional file 1: Supplementary Figures S1 to S5

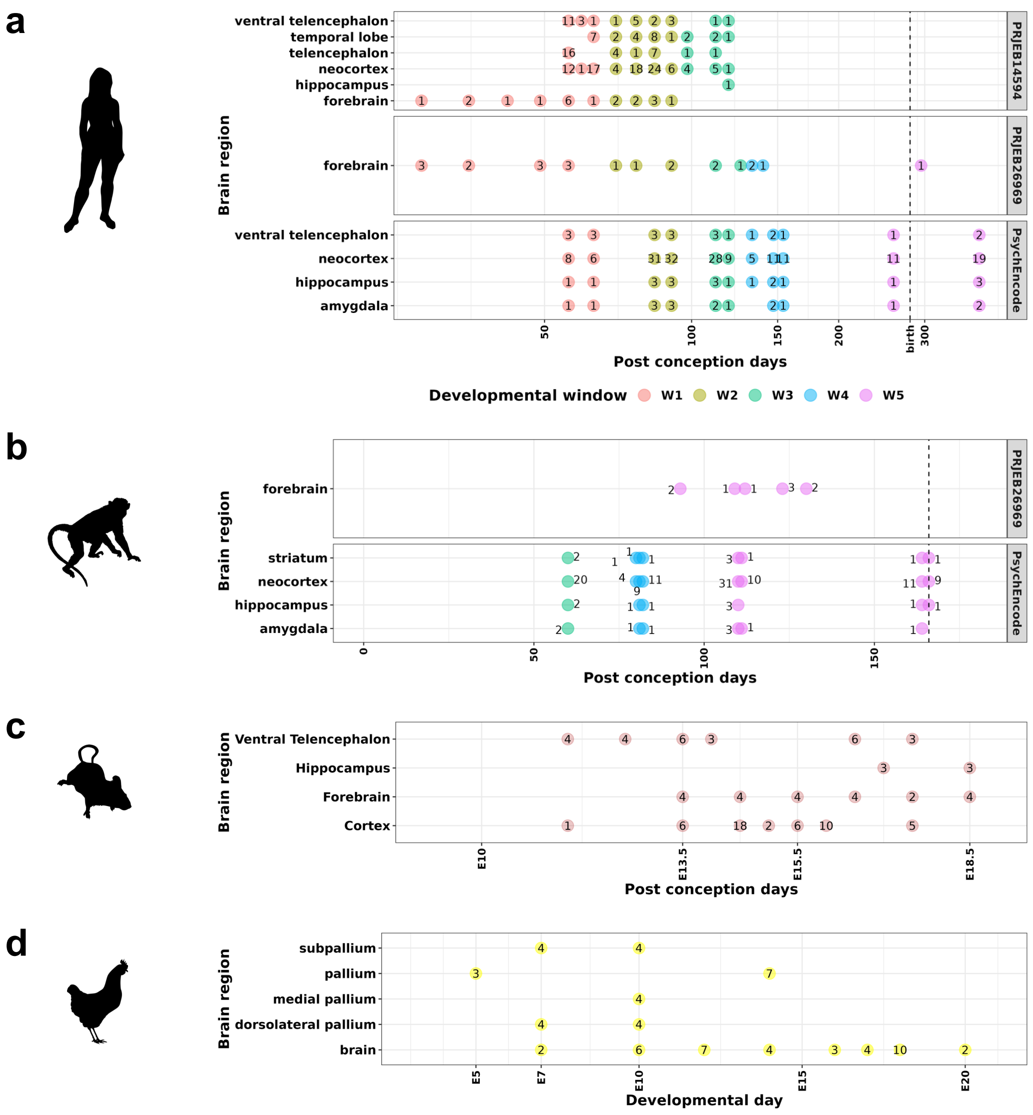


**Fig. S1. Number of samples used for annotating new transcriptome assemblies. a** Number of human samples (indicated inside the circles) used for building the transcriptome assembly of humans, grouped by the developmental time, tissue of origin, and the public data source (indicated at right), and colored by the PsychEncode developmental window. Developmental window W1, post-conception weeks 8 and 9; W2, post-conception weeks 12 and 13; W3, post-conception weeks 16 and 17; W4, post-conception weeks 19, 21 and 22; W5, post-conception week 37 and post-natal day 100. **b** Like in **a**, but for rhesus macaques. The PsychEncode developmental windows of macaques represent different developmental days but reflect similar developmental stages to humans. **c** Like in **a**, but in mice; there is no identification of matched developmental stages to humans, and samples span the cortical proliferative, early, and late neurogenesis stages. **d** Like in **a**, but in chickens; there is no identification of matched developmental stages to humans. Samples span the pallial proliferative, early, and late neurogenesis and gliogenic stages.

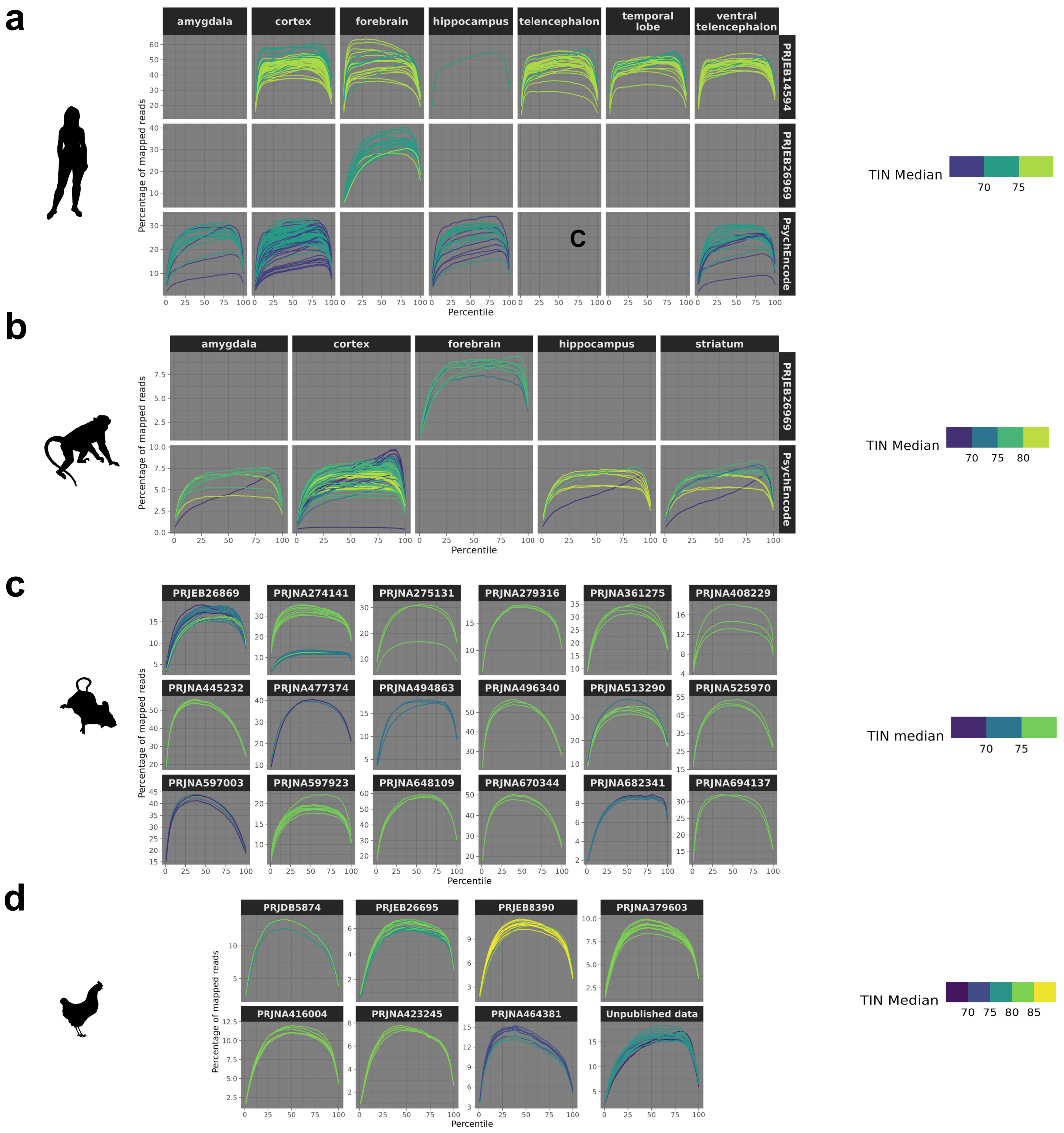


**Fig. S2. Mapping quality of samples used to generate new transcriptome assemblies.** **a** Averaged distribution of short read percentage along all transcript bodies in each library and colored depending on the Transcript Index Number (TIN) median score. Human library samples were separated by the public project and the tissues of origin. **b** Like in **a** but in the rhesus macaques. **c** Like in **a** but in mice, and libraries were separated only by the origin of public data. **d** Like in **a** but in chickens, and libraries were separated only by the origin of public data.

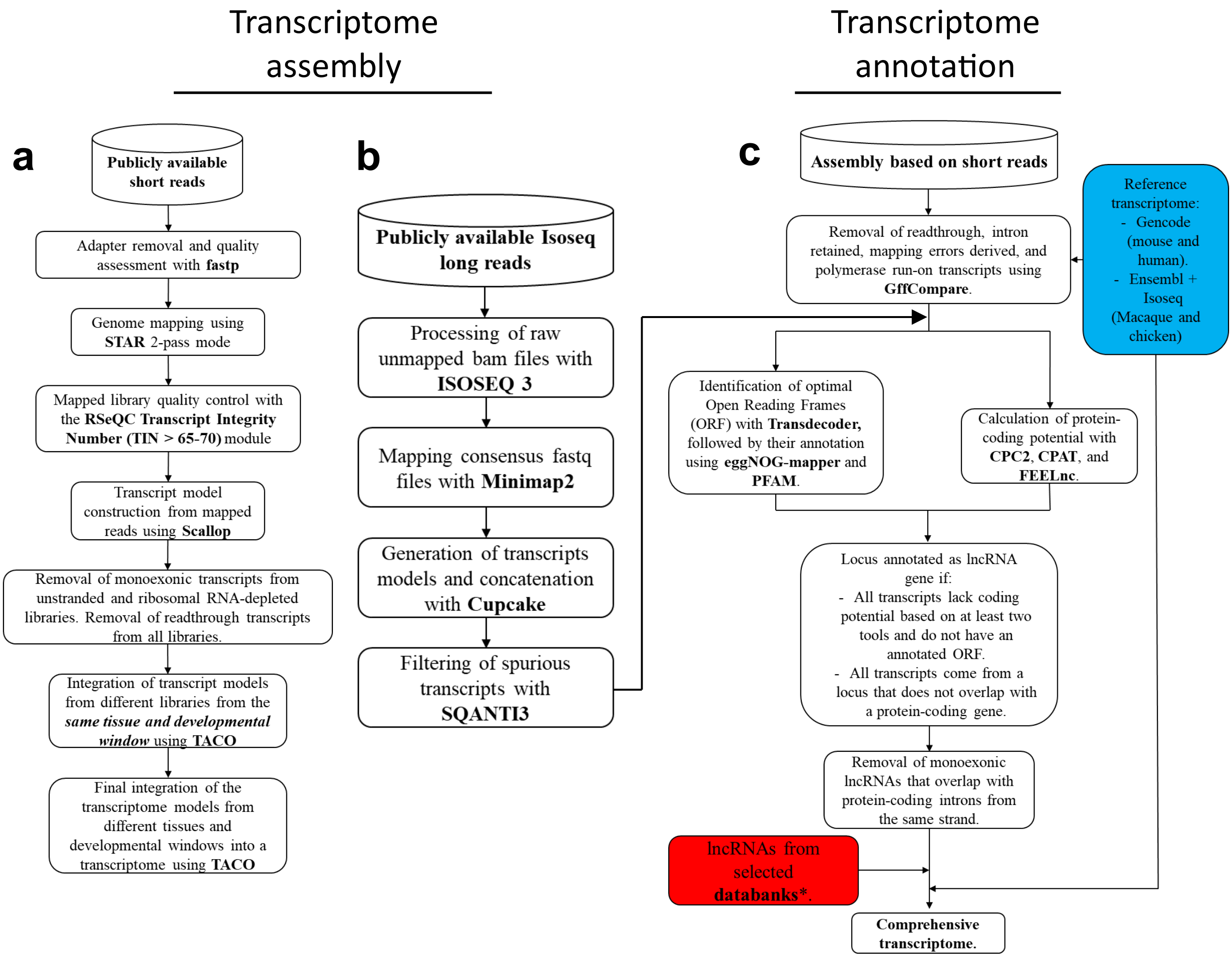


**Fig. S3. Flowcharts depicting pipelines used for identifying and annotating comprehensive transcriptomes. a** Bioinformatic pipeline developed to assemble new transcriptomes based on short reads. The pipeline uses *STAR* to map short reads, *Scallop* to build transcriptional models for each library, and *TACO* to generate consensus transcriptomes based on a set of transcriptional models. **b** Bioinformatic pipeline used to assemble new transcriptome models based on Iso-seq long reads. **c** Raw assembled transcriptomes built using short and long reads pipelines underwent extensive filters, first to remove spurious transcripts; and to identify lncRNA genes and separate them from protein-coding isoforms. Additionally, transcripts from other public databases and the lncRNA set of reference transcriptomes were added to the final comprehensive transcriptome.

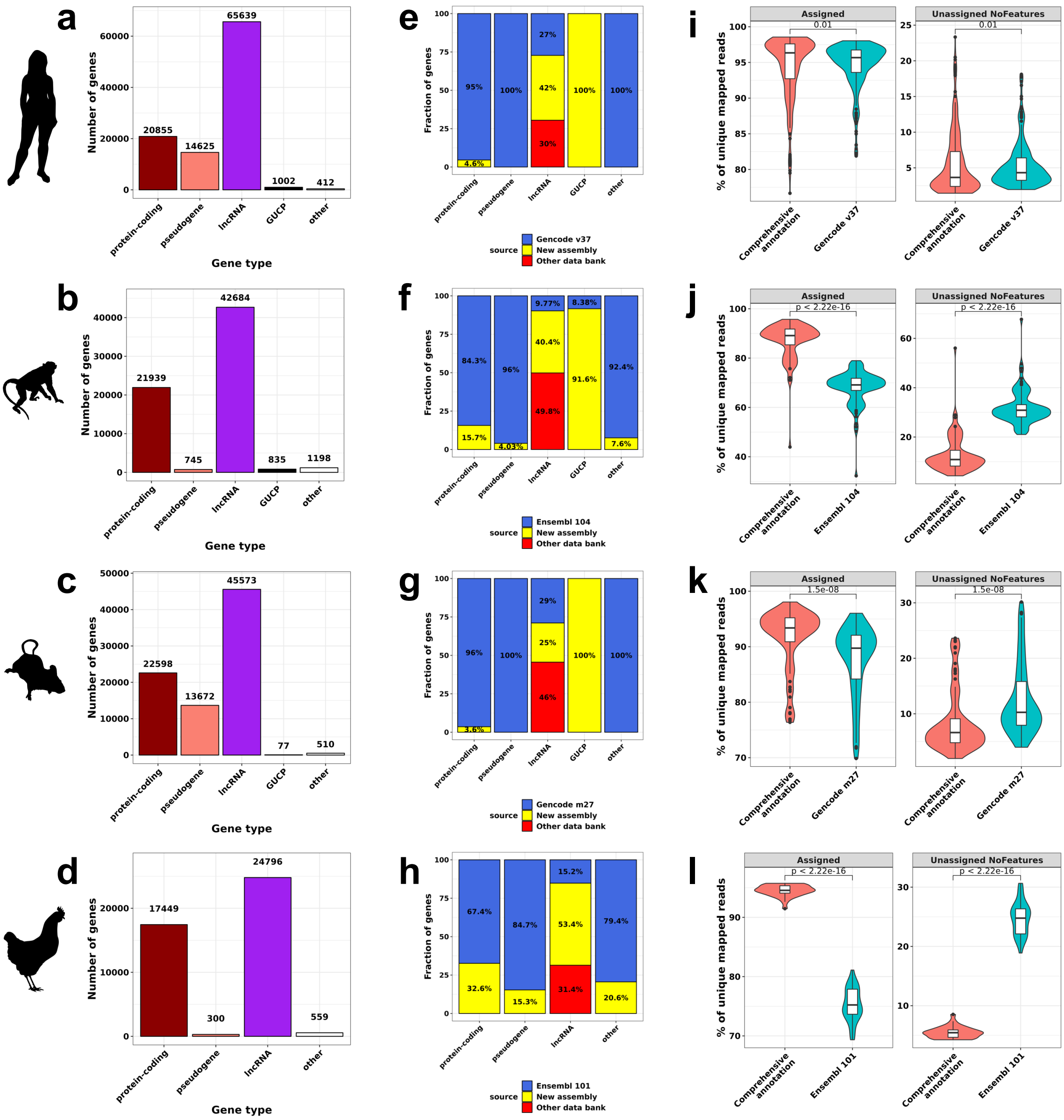

**Fig. S4. New comprehensive transcriptome assemblies improve the annotation of lncRNAs. a-d** Distribution of gene types in the new comprehensive transcriptomes annotated in the present work for humans, macaques, mice, and chickens, respectively. GUCP, genes of unknown coding potential. **e-h** Percentage of genes from different sources across the different gene types for humans, macaques, mice, and chickens, respectively. **i-l** Percentage of uniquely mapped reads, using as reference the present comprehensive annotation (red violin plots) or the Gencode and Ensembl public annotations (green violin plots), which mapped to an annotated region (Assigned, left panel) or to an unannotated region of the genome (Unassigned NoFeatures, right panel) for humans, macaques, mice, and chickens, respectively. **Statistics:** All statistics are one-sided (greater) Wilcoxon tests. ns: p greater than 0.05, *: p equal to or less than 0.05, **: p equal to or less than 0.01, ***: p equal to or less than 0.001, ****: p equal to or less than 0.0001.

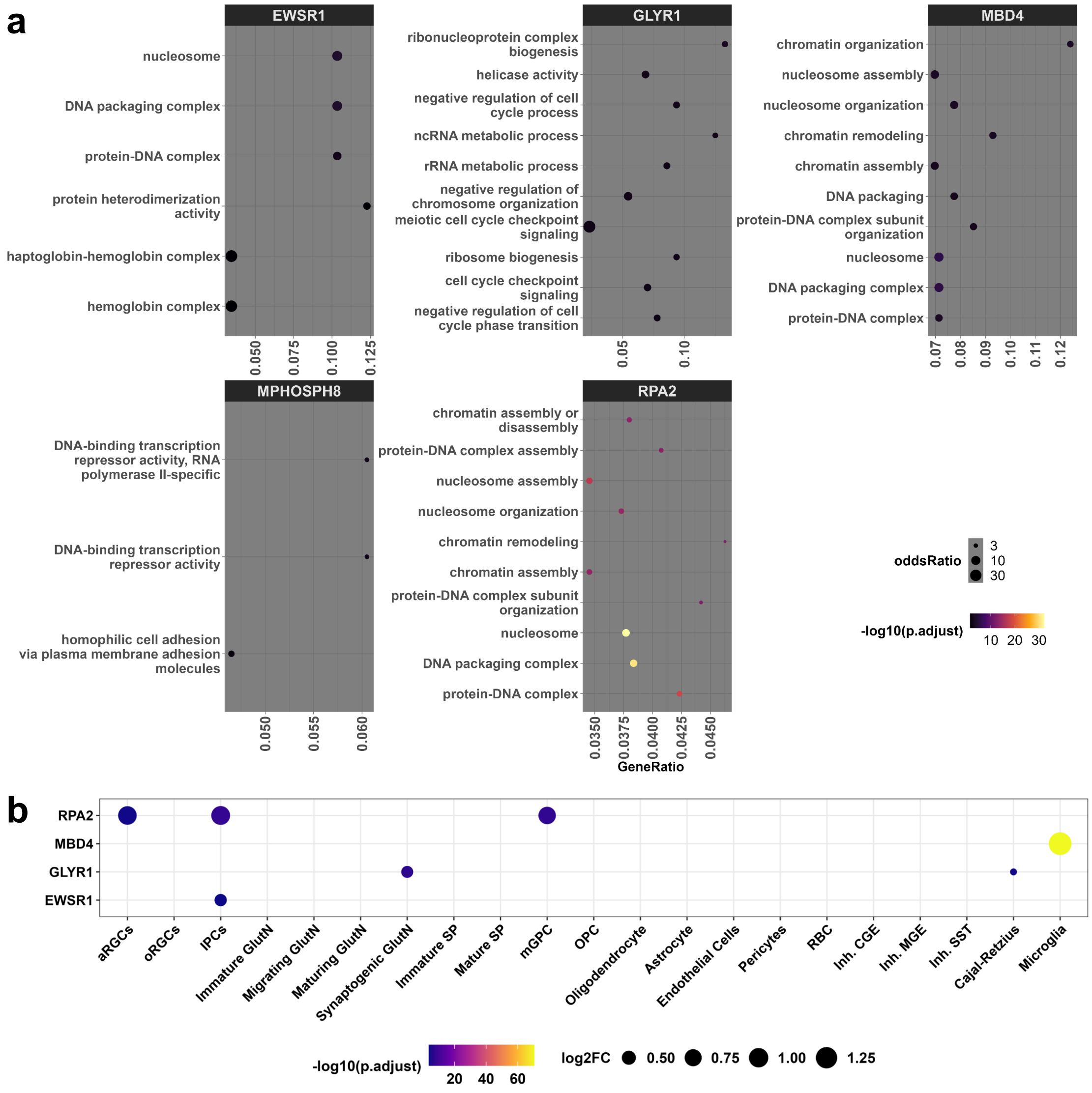


**Fig. S5. Functional features of transcription factors enriched in the promoter of young cortical lncRNAs. a** Gene ontology enrichment of genes regulated by the transcription factor (TF) indicated at the top of each panel, which is enriched in promoters of primate and Human-specific lncRNAs. **b** Dotplot showing the tissues of expression enrichment of each TF seen in **a**.
